# Actin and Src-family kinases regulate nuclear YAP1 and its export

**DOI:** 10.1101/201004

**Authors:** Nil Ege, Anna M Dowbaj, Ming Jiang, Michael Howell, Robert P Jenkins, Erik Sahai

## Abstract

The transcriptional regulator YAP1 is critical for the pathological activation of fibroblasts. In normal fibroblasts YAP1 is predominantly located in the cytoplasm, while in activated cancer-associated fibroblasts it exhibits nuclear localization and promotes the expression of many genes required for pro-tumorigenic functions. Here, we investigate the dynamics of YAP1 shuttling in normal and activated fibroblasts, using EYFP-YAP1, quantitative photo-bleaching methods, and mathematical modeling. We find that both 14-3-3 and TEAD binding modulate YAP1 shuttling, but neither affects nuclear import. Instead, we find that YAP1 serine phosphorylation is required for nuclear export. Furthermore, YAP1 nuclear accumulation in activated fibroblasts results from Src and actomyosin-dependent suppression of phosphorylated YAP1 export. Finally, we show that nuclear constrained YAP1, upon XPO1 depletion, remains sensitive to blockade of actomyosin function. Together, these data place nuclear export at the center of YAP1 regulation and indicate that the cytoskeleton can regulate YAP1 within the nucleus.

## Highlights

Photo-bleaching coupled with mathematical modeling identifies YAP1 dynamics

Regulation of nuclear export is key determinant of YAP1 localization

Serine phosphorylation is required for YAP1 nuclear export through XPO1

Nuclear YAP1 remains sensitive to actin and Src-family kinase regulation

## Introduction

The transmission of signals from the cytoplasm to transcriptional machinery in the nucleus is critical for development, tissue homeostasis, and in pathological conditions. This can occur in many ways. Signal transducing kinases can enter the nucleus and modulate transcription factor activity (Taagepera et al., 1998). Alternatively, DNA binding transcription factors can shuttle between the cytoplasm and the nucleus (Nicolás et al., 2004; Vartiainen et al., 2007; Xu and Massague, 2004). YAP1 and TAZ (WWTR1) are transcriptional regulators that are believed to be sequestered in the cytoplasm via interaction with 14-3-3 proteins when phosphorylated. In the absence of phosphorylation, YAP1 and TAZ are released and can interact with TEAD transcription factors in the nucleus (Piccolo et al., 2014; Zhao et al., 2011). Structural studies have shown that the YAP1/TEADs interaction is critically dependent on serine 94 in YAP1 (Chen et al., 2010; Li et al., 2010; Zhao et al., 2008). YAP1 and TAZ are negatively regulated by serine phosphorylation by the LATS1/2 kinases, which are themselves regulated by the MST1/2 kinases (Chan et al., 2005; Dong et al., 2007; Hao et al., 2008; Lei et al., 2008; Oka et al., 2008; Zhao et al., 2007). This pathway, often called the Hippo pathway, is critical for controlling the extent of tissue growth and organ size in *Drosophila* and mammals (Dong et al., 2007; Halder and Johnson, 2011; Pan, 2010; Zhao et al., 2011). It is regulated by a network of epithelial junctional molecules that transmit information about tissue integrity. Further, regulation by glucagon and other soluble factors couples tissue growth to nutrient availability (Enzo et al., 2015; Santinon et al., 2016; Yu et al., 2012). In all these cases, the activity of YAP1 and TAZ is negatively regulated by direct LATS1/2-mediated serine phosphorylation, including on S127 in YAP1 (Zhao et al., 2007). Thus, low levels of YAP1 and TAZ phosphorylation are linked to nuclear accumulation, leading to cell proliferation during development, wound healing, or tissue regeneration (Camargo et al., 2007; Dong et al., 2007; Gao et al., 2013; Lavado et al., 2013; Schlegelmilch et al., 2011; Zhao et al., 2011). High levels of phosphorylation are linked to cell quiescence via promotion of complexes with 14-3-3 proteins in the cytoplasm (Aitken, 2006; Muslin and Xing, 2000).

In addition to the ‘canonical’ YAP1/TAZ regulation described above, mechanical cues and tyrosine phosphorylation can modulate YAP1 function (Dupont et al., 2011; Li et al., 2016). Mechanical cues transmitted via the actin cytoskeleton are proposed to enable epithelial cells to monitor organ size (Benham-Pyle et al., 2015; Fernandez et al., 2011; Porazinski et al., 2015; Sansores-Garcia et al., 2011). This may partly depend on Src-mediated phosphorylation of tyrosine 357; however, the full details of how mechanical cues regulate YAP1 are not determined. Functional studies have shown that YAP1, but not TAZ, is critical for the pathological activation of fibroblasts in tumours and some fibrotic conditions (Calvo et al., 2013). This activation is correlated with an increased nuclear accumulation of YAP1, even though levels of S127 phosphorylation and LATS1/2 activity are not changed. The nuclear accumulation of YAP1 in cancer-associated fibroblasts (CAFs) can be reverted by perturbation of actin polymerization, actomyosin function, or tyrosine phosphorylation.

Surprisingly little is known about the dynamics of YAP1 shuttling in and out of the nucleus (Zhao et al., 2007). Many binding partners have been identified, including the TEADs, 14-3-3, and many other cytoplasmic proteins localized at cell junctions (Couzens et al., 2013; Moya and Halder, 2014), yet, it remains unclear if YAP1 is stably sequestered at these sites in either the cytoplasm or the nucleus. The rate of YAP1 shuttling between the cytoplasm and nucleus is not known, and apart from the implication of XPO1 (also called Exportin1 or Crm1) in YAP1 nuclear export (Dupont et al., 2011; Ren et al., 2010; Wei et al., 2015), the machinery regulating YAP1 entry and exit from the nucleus is not known. In this study, we answer these questions by using a variety of live imaging methods and mathematical analysis. Monitoring changes in the distribution of fluorescent proteins during or following photo-bleaching processes can provide information about protein dynamics (Nicolás et al., 2004; Vartiainen et al., 2007). Fluorescence Recovery After Photobleaching (FRAP) is used to provide information about sequestration, diffusion, and the rate of dissociation from TEAD transcription factors. Fluorescence Loss In Photobleaching (FLIP) is used to assess nuclear import and export rates, rate of TEAD association, and confirms aspects of the sequestration analysis. By combining these methods with YAP1 point mutations, actomyosin manipulations, and a screen for regulators of YAP1 nuclear import/export, we are able to derive a detailed model of YAP1 dynamics in normal fibroblasts (NFs) and pathologically activated fibroblasts (CAFs).

## Results

### Establishment of a functional YAP1 fluorescent protein

We previously reported the importance of elevated YAP1 activity in cancer-associated fibroblasts compared to normal fibroblasts (Calvo et al., 2013). This was not linked to changes in YAP1 protein levels or LATS1/2-mediated phosphorylation, but was correlated with increased nuclear localization. To image the localization and dynamics of YAP1 in normal mammary fibroblasts (NF1) and mammary carcinoma-associated fibroblasts (CAF1), we fused EYFP to the N-terminus of the protein (Figure1A). Fluorescence activated cell sorting (FACS) was used to isolate cells stably expressing low levels of EYFP-YAP1. Subsequent western blot analysis revealed that exogenous EYFP-YAP1 was expressed at just over the level of the endogenous YAP1 (FigureS1A). Further analysis with defined amounts of recombinant fluorescent protein enabled us to estimate that ∼130 000 EYFP-YAP1 molecules were expressed per cell (FigureS1B). Functional matrix contraction assays demonstrated that the expression of EYFP-YAP1 in both cells did not erroneously activate them (FigureS1C). Thus, the levels of EYFP-YAP1 expression do not greatly perturb expected function of the fibroblasts.

**Figure 1.**
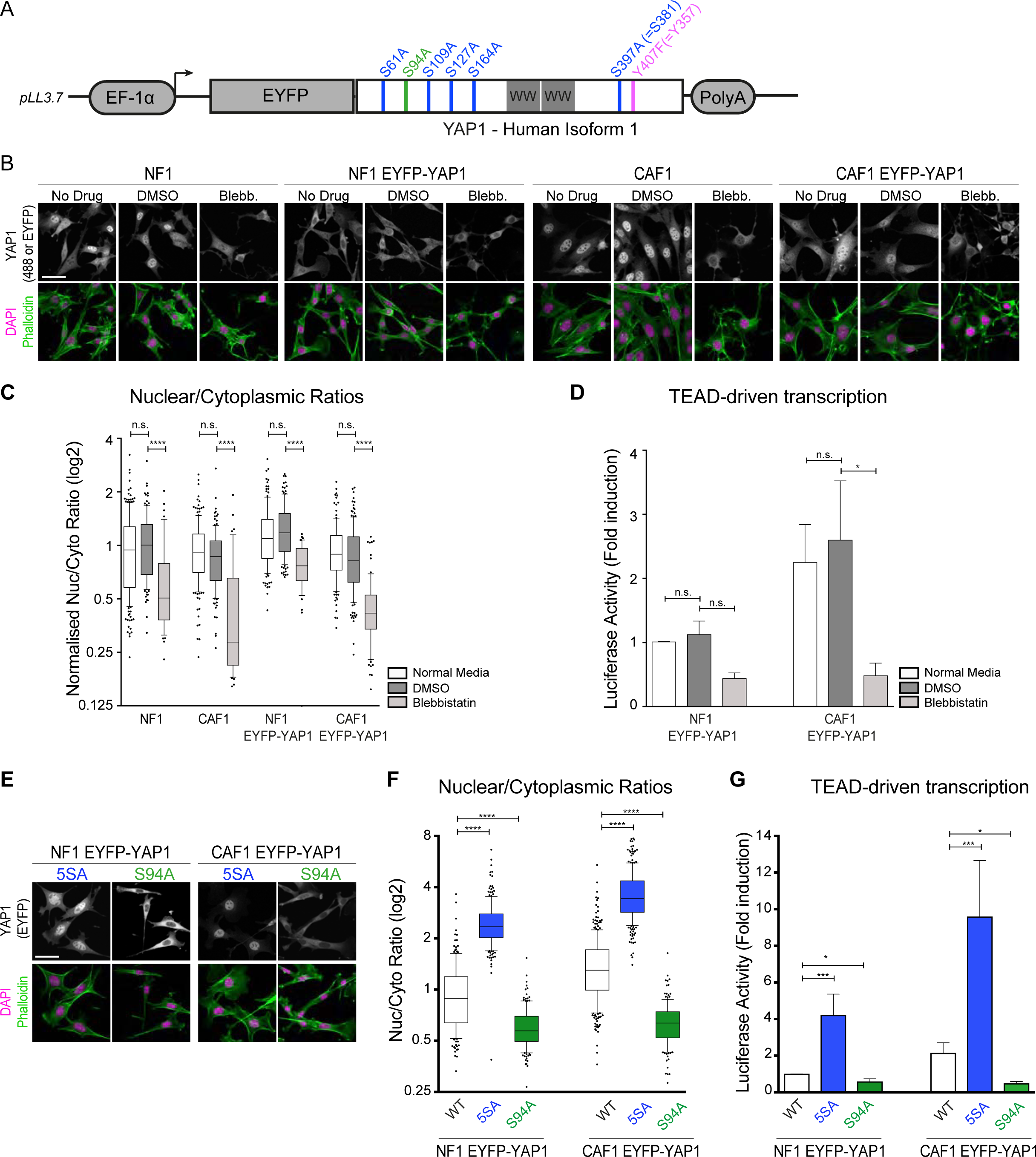
Establishment of a functional YAP1 fluorescent protein. (**A**) Schematic of the EYFP-YAP1 construct used in this study. (**B**) Representative images of endogenous YAP1 and EYFP-YAP1 localization in NF1 and CAF1 in normal condition and treated with DMSO or blebbistatin. Scale bar, 50μm. (**C**) Boxplot (with whiskers showing 10 and 90 percentiles) of nuclear-to-cytoplasmic ratio (log2 scale) corresponding to quantification of (A). n>44 cells for each condition from at least 2 independent experiments. Data are normalised to normal conditions for each cell line. (**D**) Luciferase assay of WT or EYFP-YAP1 expressing NF1 and CAF1 in normal condition and treated with DMSO or Blebbistatin. Bars represent mean ± s.e.m. of 4 independent experiments. Data are normalised to NF1 in normal media. (**E**) Representative images of EYFP-YAP1_5SA and EYFP-YAP1_S94A localization in NF1 and CAF1. Scale bar, 50μm. (**F**) Boxplot (10&90) of nuclear-to-cytoplasmic ratio (log2 scale) corresponding to quantification of (E). n>150 cells for each condition from at least 4 independent experiments. (**G**) Luciferase assay of EYFP-YAP1_5SA and EYFP-YAP1_S94A expressing NF1 and CAF1 in normal condition or treated with DMSO or Blebbistatin. Bars represent mean ± s.e.m. of 8 independent experiments. Mann-Whitney U-test, n.s., non significant, * p≤0.05**, p≤0.01, *** p≤0.001, **** p≤0.0001. See also FigureS1.

We confirmed that EYFP-YAP1 is regulated in a similar manner to endogenous YAP1 by comparing its localization in control conditions and upon blockade of actomyosin contractility. Figure1B&C shows that EYFP-YAP1 is more nuclear in CAF1 than NF1; this mirrors the difference in endogenous protein localization (FigureS1D). More significantly, EYFP-YAP1 showed a similar cytoplasmic shift to the endogenous protein upon actomyosin blockade using blebbistatin. These results were also observed at the transcriptional level in CAF1 (Figure1D). No statistical difference was observed in NF1 upon blebbistatin treatment, probably due to the already low level of transcription before treatment. We further probed the behavior of the EYFP-YAP1 fusion by introducing well-characterized mutations at serine 94 (S94A) (Zhao et al., 2008) and serines 61, 109, 127, 164 and 381 (termed 5SA) (Zhao et al., 2007), blocking the interaction of YAP1 with TEAD and 14-3-3, respectively (Figure1A and FigureS1E). Both mutants showed the expected cytoplasmic and nuclear localization, respectively (Figure1E&F). The functionality of the EYFP-YAP1 was also evidenced by the increased TEAD reporter activity in cells expressing EYFP-YAP1_5SA (Figure1G). Cells expressing EYFP-YAP1_S94A showed reduced TEAD reporter and matrix contraction activity indicating that this construct acts as a dominant negative (Figure1G and FigureS1F).

### YAP1 is not stably sequestered in the cytoplasm or the nucleus

Having established that EYFP-YAP1 is a valid tool to probe YAP1 function, we embarked on Fluorescence Recovery After Photobleaching (FRAP) experiments combined with mathematical modeling in order to assess protein diffusion (D) and dissociation rates (k_-0_ and k_-1_) (Figure2A) (Fritzsche and Charras, 2015; Sprague et al., 2004). During FRAP, a region of interest was bleached and the time for fluorescence recovery compared to an adjacent non-bleached region was assessed. An incomplete recovery would indicate an ‘immobile fraction’ sequestered in the compartment of interest. FRAP analyses in both the cytoplasm and the nucleus of NF1 and CAF1 were performed. Figure2B shows that the bleached area has a fluorescent intensity equivalent to a non-bleached region within ∼15s (See also FigureS2A). This indicates that there is no ‘immobile’ fraction of EYFP-YAP1 in the time-frame of our experiments and thus that the molecule does not engage in permanent binding to a fixed component in the nucleus or in the cytoplasm. We confirmed the validity of our FRAP experiments on fast-diffusing EGFP and stably chromatin-bound H2B-GFP (FigureS2B&C and Movie1). The H2B-GFP analysis also demonstrates that chromatin can be considered immobile in the time-frame of our experiments.

### Binding to TEAD transcription factors modulates YAP1 nuclear mobility

Both diffusion of EYFP-YAP1 and its release from a short-lasting interaction could influence the observed rate of fluorescence recovery (half-time), the effective radius (r_e_) and the depth (K) of the post-bleach profile (FigureS2D and Mathematical Methods) (Fritzsche and Charras, 2015; Kang et al., 2010; Kang et al., 2009). If diffusion is slow enough, these parameters would change according to the size of the bleached region as diffusion into a larger region will take longer (FigureS2D-Situation1). We therefore repeated FRAP analyses with different sized bleached regions in the nucleus and the cytoplasm. The intensity recoveries (Figure2B) and the post-bleach profile (FigureS3A) showed no differences between the three bleached regions. Furthermore, the half-time, the effective radius and the bleach-depth did not change significantly (Figure2C and FigureS3). Taken together, these results indicate that diffusion is rapid relative to the imaging rate of 16.7Hz suggesting that the recovery observed reflects unbinding/binding reactions (FigureS2D–Situation2). Consistent with this, Figure2D shows that a pure diffusion model did not fit the experimental recovery curves well (orange vs blue curves). Both reaction-diffusion (red) and reaction models (green and magenta) fitted the data well. Akaike Information Criterion (AIC) analysis indicated that in the majority of cases reaction-based models fitted best (Figure2G, TableS1 and Mathematical Methods). In cases where reaction-diffusion models fitted well, the diffusion value was in the range of 25-40 μm^2^s^-1^, which is in the range reported for multimerized GFP with similar molecular mass to EYFP-YAP1 (Baum et al., 2014). Therefore, diffusion is so rapid that it has largely occurred by the first post-bleach image acquisition and the subsequent recovery captured in our analyses mainly reflects the reaction component, such as the dissociation of molecules from their bound immobile state (Figure2E&F). Although a two-reactions model fits best to a third of CAF1 data (Figure2G), it did not improve the match to the experimental data enough to justify the increase in number of parameters (TableS1). We therefore discarded this approach as unnecessarily complex and used a one-reaction model to extract YAP1 dissociation rate. Intriguingly, this rate in the nucleus was significantly lower in CAF1 (0.39s^-1^) than in NF1 (0.56s^-1^) (Figure2H and Movie2). This suggests that YAP1 associates more stably with a nuclear partner in CAFs, likely a chromatin bound factor. Similarly, FRAP analyses in the cytoplasm determined that the observed fluorescence recoveries could be explained by dissociation rates of 0.67s^-1^ in NF1 and 0.65s^-1^ in CAF1 (Figure2H and Movie3). This means that there is relatively little difference between normal and activated fibroblasts and that YAP1 does not have a long-lived site of sequestration in the cytoplasm.

**Figure 2:**
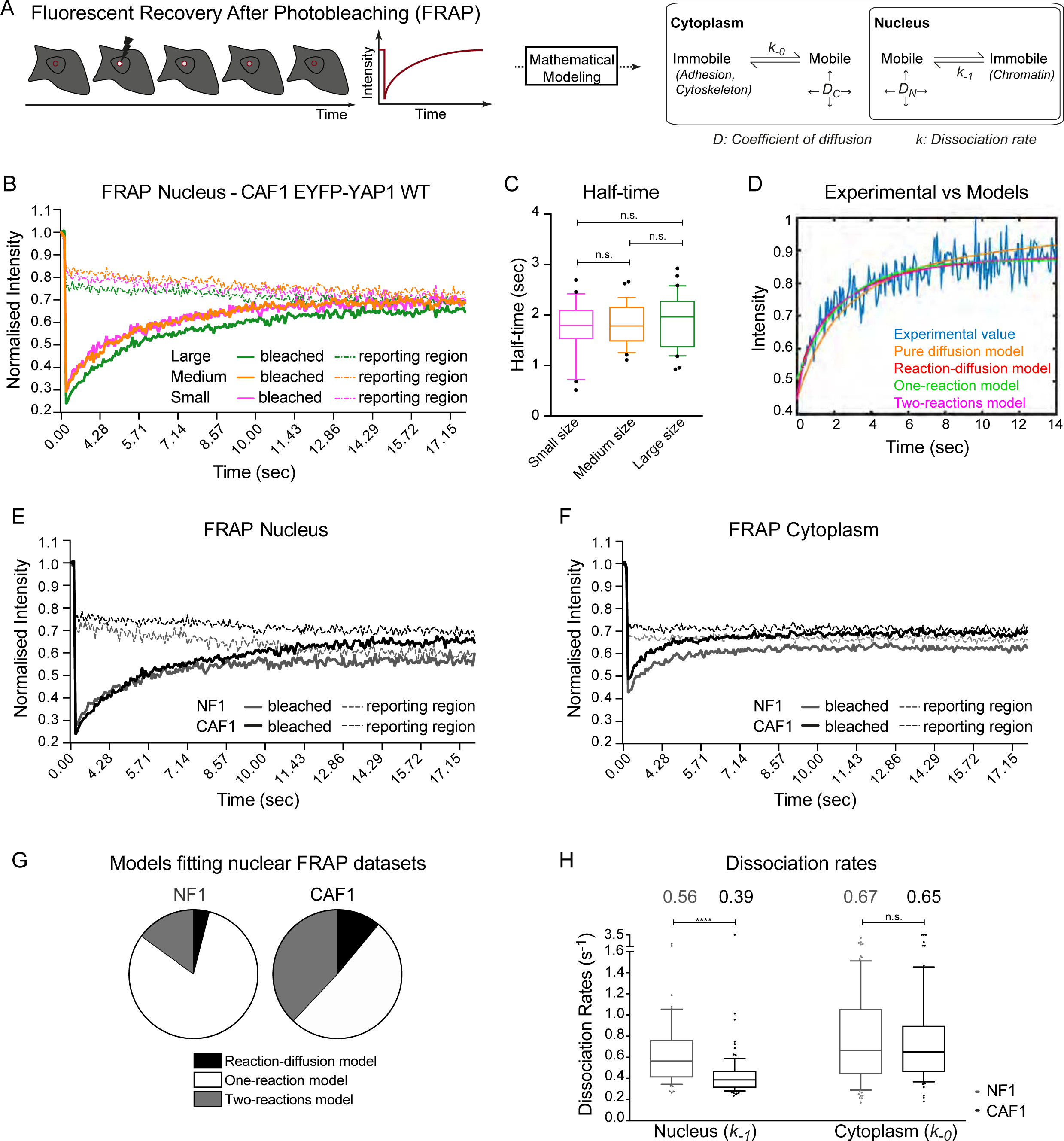
YAP1 is not stably sequestered in the cytoplasm or the nucleus. (**A**) Schematic of the FRAP experiment and parameters extracted with mathematical modeling. (**B**) Graph showing the median of EYFP-YAP1 intensities of three different sized bleached (plain line) and reporting (dotted line) regions during nuclear FRAP in CAF1. n > 29 cells for each sizes. (**C**) Boxplot (10&90) of half-time corresponding to recovery curves from (B).(**D**) One representative example of the fits of four models to experimental values. (**E**) Equivalent graph to (B) upon nuclear FRAP in NF1 (n=20 cells) and CAF1 (n=30 cells), from 3 biological replicates. (**F**) Equivalent graph to (B) upon cytoplasmic FRAP in NF1 (n=30 cells) and CAF1 (n=30 cells), from 3 biological replicates. (**G**) Pie charts showing the distribution in percentage of best fitting model for nuclear FRAP in NF1 and CAF1. (**H**) Box-plot (10&90) showing the dissociation rates of EYFP-YAP1 in the nucleus and the cytoplasm of NF1 and CAF1 from 3 biological replicates. n=59(nuc)/81(cyto) cells for NF1 and n=88(nuc)/72(cyto) cells for CAF1. Values above plots indicate medians. Noisy cells are assumed to have rapidly recovered and are represented with a large arbitrary unit of 3.5s^-1^. Mann-Whitney U-test, n.s., non significant, **** p<0.0001. See also FigureS2 and S3.

Having established that YAP1 has differential binding in the nucleus in CAF1 compared to NF1, we sought to determine what the binding partner was. The most likely partners of YAP1 in the nucleus are TEAD transcription factors (Zanconato et al., 2015; Zhao et al., 2008). We therefore repeated the FRAP analyses with S94A mutation, which is known to abrogate TEAD interaction. Figure3A&B show that S94A mutation did indeed affect fluorescence recovery (See also Movie4). In some cases, it became so rapid that it was not possible to reliably determine a dissociation rate. In the CAFs in which a rate could be measured, it was 0.97s^-1^ while the median rate of the whole data set was 1.29s^-1^ (Figure3E). These data confirm that the altered dynamics of EYFP-YAP1 in the nucleus of CAF1 is due to TEAD binding. In contrast, the active 5SA mutant, unable to bind 14-3-3 proteins, exhibited a slower dissociation rate, consistent with increased TEAD binding and transcriptional activation (Figure3C-E and Movie4). To determine if the dissociation rate measured in CAF1 (0.21s^-1^) represents the dissociation rate of an intact YAP1-TEAD complex from chromatin or the dissociation rate of YAP1 from TEAD that remains bound to chromatin, we performed FRAP of TEAD1-mCherry. Figure3F&G show that TEAD1-mCherry had a slower recovery than YAP1 with a dissociation rate of 0.05/0.06s^-1^ (See also Movie5). This suggests that in CAFs, TEAD1 typically spends 30s bound to chromatin whereas YAP1 typically spends ∼2.5s bound to TEADs, indicating that the rate we measured represents the dissociation of YAP1 from TEAD transcription factors.

**Figure 3:**
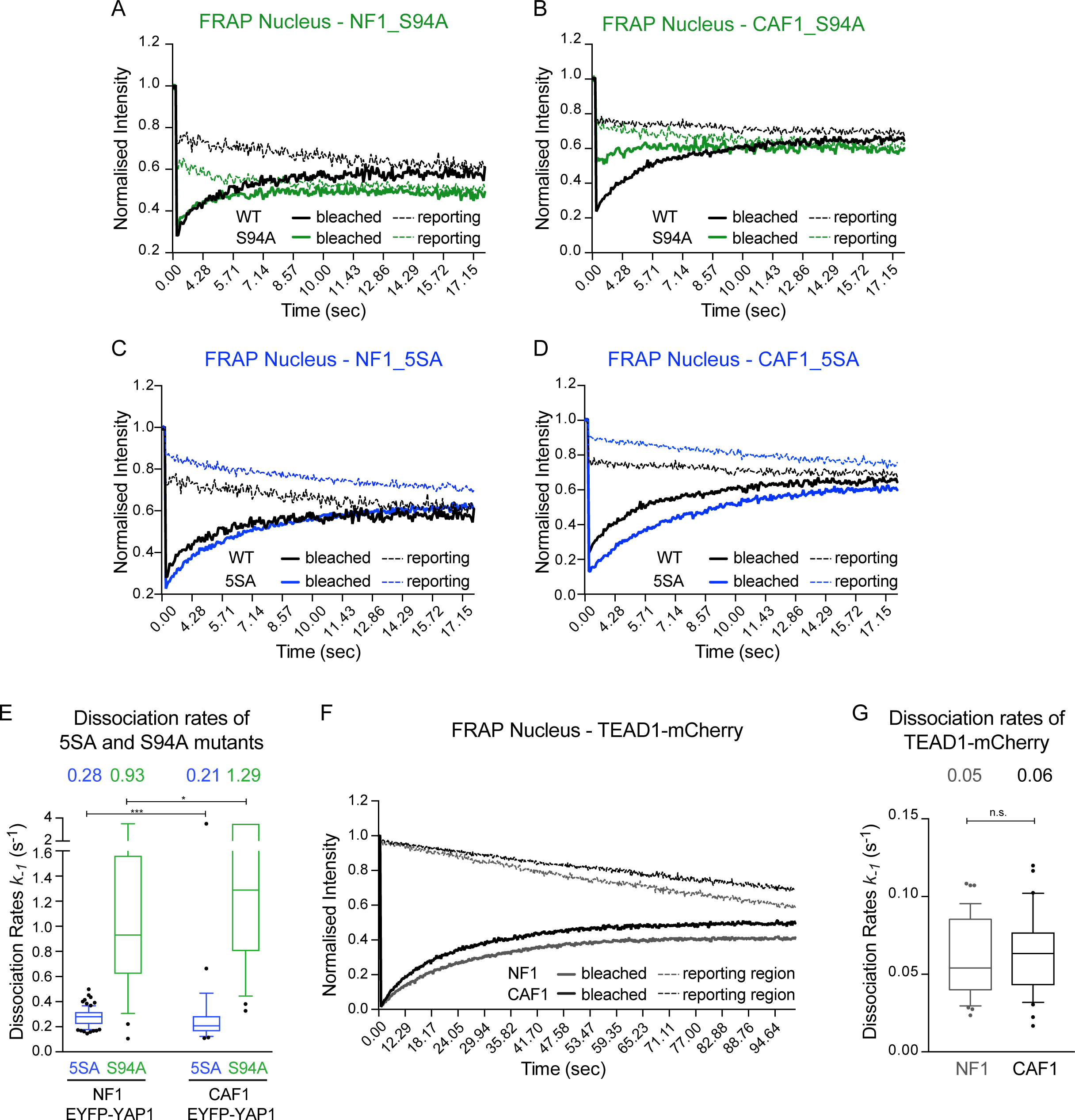
Binding to TEAD transcription factors modulates YAP1 nuclear mobility. (**A**) Graph showing the median intensities of EYFP-YAP1_S94A and EYFP-YAP1 (reproduced from Fig2E for representation) from bleached (plain line) and reporting (dotted line) regions upon nuclear FRAP in NF1. n=30 cells, 3 biological replicates. (**B**) Equivalent graph to (A) for nuclear FRAP in CAF1 with EYFP-YAP1_S94A, n=30 cells, 3 biological replicates. (**C**) Equivalent graph to (A) for nuclear FRAP in NF1 with EYFP-YAP1_5SA, n=30 cells, 3 biological replicates. (**D**) Equivalent graph to (A) for nuclear FRAP in CAF1 with EYFP-YAP1_5SA, n=25 cells, 3 biological replicates. (**E**) Box-plot (10&90) showing the dissociation rates of 5SA and S94A mutants in the nucleus of NF1 and CAF1. Values above plots indicate medians. Noisy cells are assumed to have rapidly recovered and are represented with a large arbitrary unit of 3.5s^-1^. (**F**) Equivalent graph to (A) for nuclear FRAP of TEAD1-mCherry in NF1 and CAF1, n>37 for both cell lines, from 3 biological replicates. (**G**) Histogram 10-90 percentiles showing the dissociation rates of TEAD1-mCherry in the nucleus of NF1 and CAF1. Values above plots indicate medians. Mann-Whitney U-test, n.s., non significant, *p≤0.05, *** p≤0.001.

### YAP1 dynamics is regulated by nuclear export

Next, we turned our attention to determining the rate of nuclear import and export. We employed Fluorescence Loss In Photobleaching (FLIP) analysis to continually bleach EYFP-YAP1 in the nucleus and assess import rates (k_-2_), export rates (k_2_) as well as the nuclear association rates (k_1_) (Figure4A). Figure4B shows the different loss of signal at the bleached point (black) and ‘reporting’ points in the nucleus (green) and the cytoplasm (orange) that were selected for being a similar distance from the bleached point. FLIP analyses on fast-diffusing EGFP revealed no differences in the loss of signals between the bleached and the reporting point in the nucleus and a fast drop of the cytoplasmic intensity (FigureS4A). The greater difference in curves of the nuclear and cytoplasmic ‘reporting’ points (green and orange) in CAF1 compared to NF1 provide a qualitative indication of reduced exchange between the nucleus and cytoplasm in these cells (Figure2B). However, we sought quantitative determination of the rates (Ungricht et al., 2015; Wustner et al., 2012). Closer inspection of the imaging data revealed that EYFP-YAP1 signal did not diminish uniformly across the nucleus (Movie6). This is not unexpected given that YAP1 dynamics in the nucleus result from the combined action of diffusive and reactive processes (i.e. TEAD binding), but this meant it would not be sensible to consider nuclear EYFP-YAP1 signal as a single variable as shown in Figure4B. To overcome this, we divided the entire cell into regions of 2.1μm^2^ and measured EYFP-YAP1 signal in each region and constructed a Partial Differential Equation (PDE) based model of YAP1 dynamics (FigureS4 and Mathematical Methods). This accounted for the mobility of YAP1 in the nucleus and in the cytoplasm. The diffusion value (*D*) and dissociation values (*k_-1_*) determined by FRAP analysis were then used to model the non-uniform decrease in EYFP-YAP1 signal across the nucleus and the import (*k_-2_*) and export (*k_2_*) rates were calculated based on EYFP-YAP1 signal intensity either side of the nuclear boundary (FigureS4B). This technique allows the model to fit the data according to all spatial locations, as opposed to just the bleached region and two user-selected regions in the nucleus and the cytoplasm, generating greater robustness in the model and certainty in the estimated parameters (FigureS4C). In addition to the import and export rates, this analysis also generated a value for the association rate (*k_1_*) of YAP1 with TEAD – that is the reverse process of the dissociation rate measured in Figure3.

**Figure 4:**
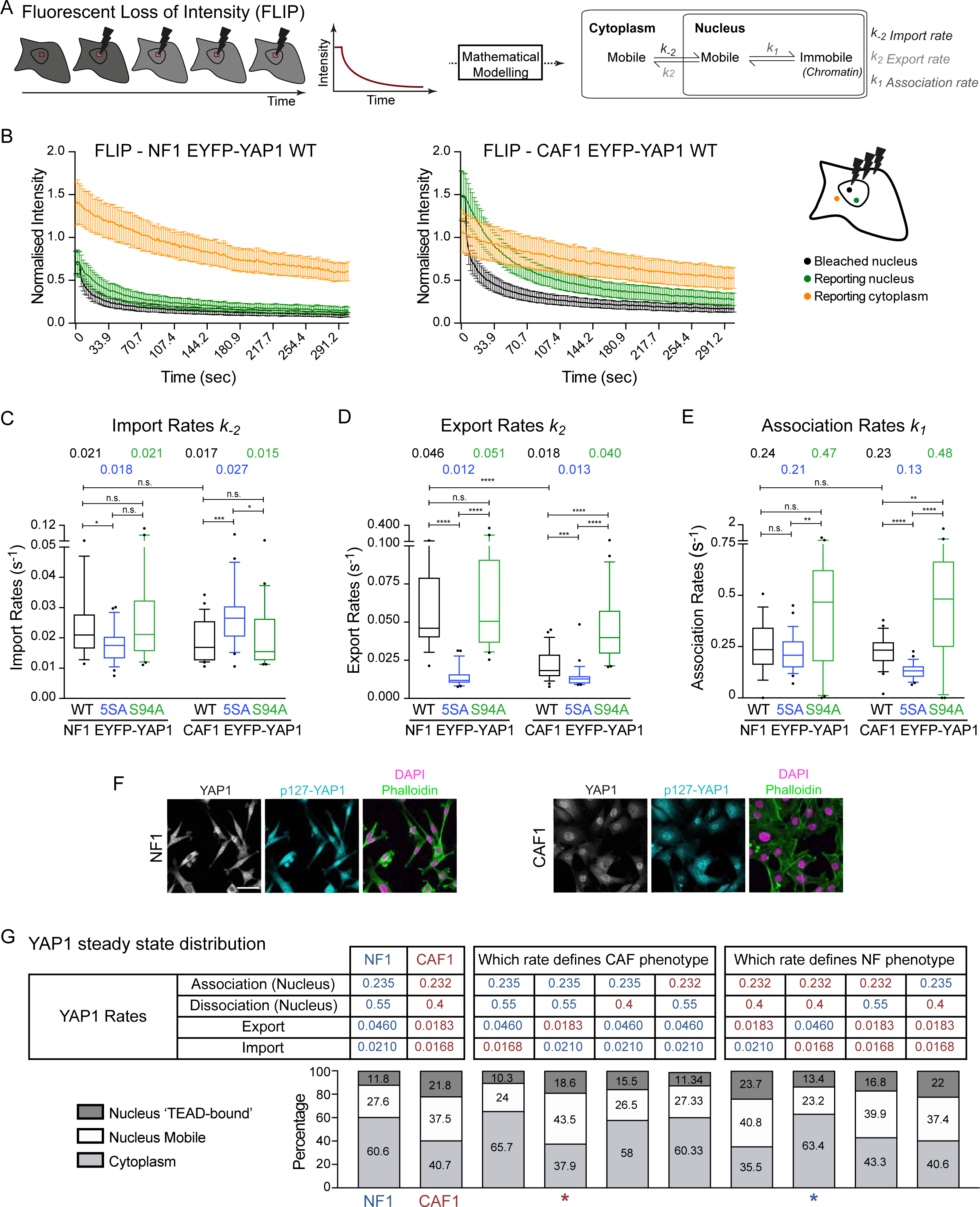
YAP1 dynamics is regulated by nuclear export. (**A**) Schematic of the FLIP experiment and parameters extracted with mathematical modeling. (**B**) Graph showing the intensities of EYFP-YAP1 WT from bleached (black), nuclear reporting (green) and cytoplasmic reporting (orange) regions upon nuclear FLIP in NF1 and CAF1. Graph represents mean with 95%CI. n=26 cells for each, from 3 biological replicates. (**C-E**) Box-plot (10&90) showing the import (*k-_2_*), export (*k_2_*) and association rates (*k_1_*) of EYFP-YAP1 WT, 5SA or S94A in NF1 and CAF1. Values above plots indicate medians. Noisy cells are assumed to have rapidly recovered and are represented with a large arbitrary unit of 3.5s^-1^. (**F**) Representative images of S127YAP1 compared to total YAP1 in NF1 and CAF1. (**G**) Estimation of YAP1 steady-state distribution in normal condition and upon modification of specific rates. Stars highlight CAF-like (red) or NF-like (blue) distributions. Mann-Whitney U-test, n.s., non significant, * p≤0.05, ** p≤0.01, *** p≤0.001, **** p≤0.0001. See also FigureS4.

The quantitative analysis described above revealed that YAP1 import was remarkably comparable between NF1 (0.021s^-1^) and CAF1 (0.017s^-1^) (Figure4C). Further, mutation of the serine residues involved in 14-3-3 binding did not greatly affect YAP1 import in NF1 (WT 0.021s^-1^ and 5SA 0.018s^-1^) (Figure4C & FigureS4D-G and Movie7). This argues against 14-3-3 mediated sequestration in the cytoplasm preventing nuclear import. Further, the increased levels of nuclear YAP1 in CAF1 cannot be attributed to faster import. Figure4D shows that the export of rate of EYFP-YAP1 was different between NF1 and CAF1. YAP1 export was 2.5 times slower in CAF1, compared to NF1 (NF1 0.046s^-1^, CAF1 0.018s^-1^). The rate of YAP1 export was also diminished by mutation of LATS1/2 target sites in both cells (NF1_5SA 0.0.012s-1s^-1^, CAF1_5SA 0.013s^-1^). However, the reduced export rate of YAP1 in CAF1 cannot be attributed to reduced LATS1/2 mediated phosphorylation as both pS127 YAP1 and active pLATS1/2 levels are equivalent between NF1 and CAF1 (FigureS4H). The slow export rates of EYFP-YAP1_5SA compared to EYFP-YAP1 further support the idea of a LATS1/2 independent mechanism of regulating export. Sensitivity analysis revealed that increasing or decreasing the dissociation rate by 50% (which approximates to the range of values measured by FRAP in Figure2) had minimal effect on the import and export rates (Mathematical Methods), increasing our confidence in our results. Together, these data establish the regulation of nuclear export as a key difference between normal and activated fibroblasts. They also suggest that nuclear export can be modulated in two ways, one dependent on and the other independent of LATS1/2 phosphorylation. The reduced export of EYFP-YAP1_5SA implies that phosphorylated YAP1 is present in the nucleus, which is contrary to the current view of its retention in the cytoplasm bound to 14-3-3 proteins. We sought direct evidence of this by performing staining of pS127-YAP1. Figure4F shows that phosphorylated YAP1 could indeed be found in the nucleus or both NF1 and CAF1.

The analysis of the FLIP data also enabled the association rate of YAP1 to TEAD to be determined. This parameter, alongside dissociation and diffusion rates, affects the spatial variability of intensity within the nucleus (Mathematical Methods). For both NF1 and CAF1, and for the various YAP1 mutants the association rate was consistently in the range of 0.15–0.5s^-1^(Figure4E). Unsurprisingly, the association rate showed greater sensitivity to changes in the dissociation rate (Mathematical Methods). Together these data now enable a description of the steady state distribution and dynamics of YAP1 in both normal and activated mouse mammary fibroblasts (Figure4G). Briefly, we determine that ∼40% of EYFP-YAP1 is in the nucleus in NF1, with ∼12% in a chromatin-bound fraction. In contrast, ∼60% of EYFP-YAP1 is nuclear in CAF1, with ∼22% bound to chromatin. This ∼50% increase in chromatin binding in CAFs is similar to the two times increase in transcriptional activity (Figure1D). Similar steady state distributions were generated for the 5SA and S94A mutants (FigureS4I). 5SA mutant distributions are similar between NF1 and CAF1, showing an increase of both nuclear fractions (chromatin bound and mobile). The steady state distributions of S94A mutant are also similar between both NF1 and CAF1 and resembled the steady state of YAP1 distribution in NF1 (See also Figure4G). Individually substituting dynamic parameters between NF1 and CAF1 indicated that simply swapping the export rate and keeping all the other NF1 parameters (red star) was sufficient to yield a CAF-like distribution of YAP1, and vice-versa (blue star) (compare columns 1, 2, 4, and 8 in Figure4G).

### Actin and Src-family kinases regulate YAP1 export

We next focused on how non-canonical regulation of YAP1 by the actomyosin cytoskeleton modulated YAP1 dynamics. The fluorescent properties of blebbistatin precluded its use in photobleaching experiments, we therefore used latrunculin B to disrupt the actin cytoskeleton (Kolega, 2004). We additionally used the Abl and Src-family kinase inhibitor dasatinib as previous analyses had implicated these kinases in ‘mechano-‘ regulation of YAP1 downstream of actomyosin contractility (Calvo et al., 2013; Elbediwy et al., 2016; Li et al., 2016; Rosenbluh et al., 2012). In agreement with previously published data, we confirmed that treatment of fibroblasts with those inhibitors induces YAP1 cytoplasmic translocation (Figure5A&B). Similarly, Figure5A&B show that both inhibitors treatments lead to EYFP-YAP1 translocation to the cytoplasm. The use of low concentrations of those inhibitors, which affect the localization without decreasing dramatically the cell size, is beneficial for photobleaching experiments. Tyrosine phosphorylation of YAP1 has been implicated as a regulatory mechanism in both epithelial cells and CAFs (Calvo et al., 2013; Levy et al., 2008; Li et al., 2016; Rosenbluh et al., 2012; Tamm et al., 2011; Zaidi et al., 2004). Therefore, we investigated whether mutation of Y357, which is a Src-family kinase and Abl phosphorylation site, would phenocopy the effect of dasatinib treatment. We confirmed that this mutation reduced the transcriptional competence of YAP1 (Figure5C) leading to a decrease of contraction ability (FigureS5C&D). However, Y357F mutant exhibited surprisingly similar sub-cellular distribution as EYFP-YAP1 (FigureS5E&F). Interestingly, steady-state distribution of Y357F mutants shows a lower fraction of YAP1 bound to the chromatin (∼10%) compared to EYFP-YAP1 (∼22%) (Figure5I). Further, FRAP and FLIP analyses confirm that mutation of Y357F had little effect on the rates of association, dissociation, import, or export (Figure5H and FigureS5G&H). These data demonstrate that the effect of dasatinib treatment cannot be accounted for by direct YAP1 tyrosine phosphorylation on Y357.

**Figure 5:**
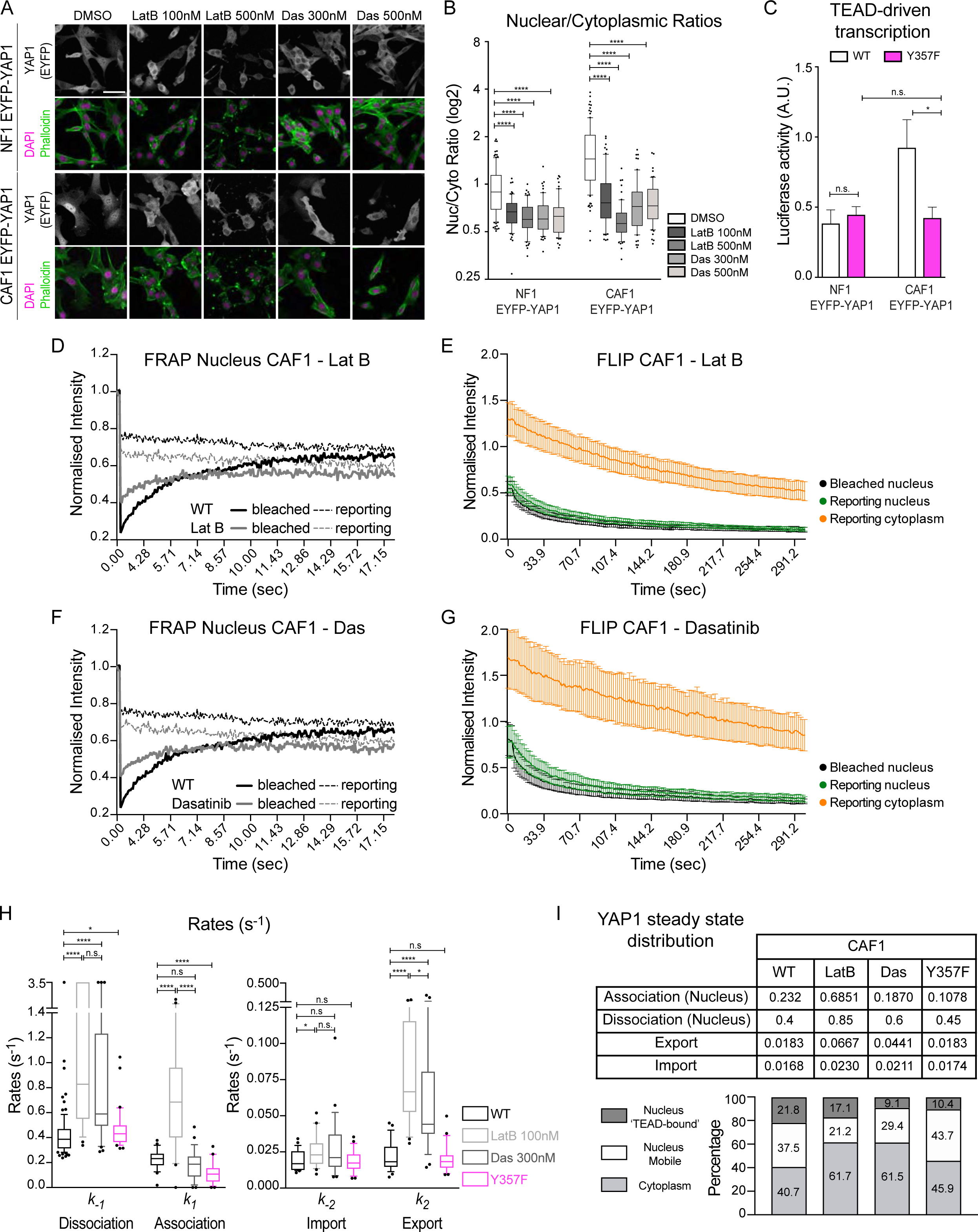
Actin and Src-family kinases regulate YAP1 export. (**A**) Representative images of EYFP-YAP1 localization in NF1 and CAF1 treated with DMSO or latrunculin B and dasatinib. Scale bar, 50μm. (**B**) Box-plot (10&90) of nuclear-to-cytoplasmic ratio (log2 scale) corresponding to (A). n>60 cells, at least 2 independent experiments. (**C**) Luciferase assay of EYFP-YAP1 vs EYFP-YAP1_Y357F in NF1 and CAF1. Bars represent mean ± s.e.m. of at least 6 independent experiments. (**D**) Graph showing the median intensities of EYFP-YAP1 from bleached (plain line) and reporting (dotted line) regions upon nuclear FRAP in CAF1 treated with latrunculin B, n= 29 cells, 3 biological replicates. EYFP-YAP1 WT with no treatment is reproduced from Fig2E for representation. (**E**) Graph showing the intensities of EYFP-YAP1 from bleached (black), nuclear reporting (green) and cytoplasmic reporting (orange) regions upon nuclear FLIP in CAF1 treated with latrunculin B. Graph represents mean with 95%CI. n= 30 cells, 3 biological replicates. (**F**) Equivalent graph to (D) upon nuclear FRAP in CAF1 treated with dasatinib, n= 30 cells, 3 biological replicates. EYFP-YAP1 WT with no treatment is reproduced from Fig2E for representation. (**G**) Equivalent to (E) upon FLIP in CAF1 treated with dasatinib, n=30 cells, 3 biological replicates. (**H**) Box-plot (10&90) showing the different rates. Rates of EYFP-YAP1 WT are reproduced from Fig2h for representation. Noisy cells are assumed to have rapidly recovered and are represented with a large arbitrary unit of 3.5s^-1^. Medians are indicated in the table (I). (**I**) Estimation of YAP1 steady-state distribution. EYFP-YA1 WT in CAF1 is reproduced from Fig4G for representation Mann-Whitney U-test, n.s., non significant *, p≤0.05, ** p≤0.01, *** p≤0.001, **** p≤0.0001. See also Figure S5.

FRAP and FLIP analysis revealed that both latrunculin B and dasatinib increased the dissociation rate of YAP1 from chromatin (Figure5D-H). The association rate of YAP1 was increased after latrunculin B treatment, but not after dasatinib treatment. More crucially, YAP1 export rates increased significantly following latrunculin B and dasatinib treatment (Figure 5H), returning to rates similar to those in NF1 (Figure3H). Furthermore, EYFP-YAP1 CAF1 treated with latrunculin B or dasatinib, present similar protein distribution that in NF1 (Figure5I compared to Figure4G). S127 phosphorylation of YAP1 was not affected by latrunculin B or dasatinib treatment (FigureS5I), however, the non-phosphorylated EYFP-YAP1_5SA mutant remained nuclear even in latrunculin B and dasatinib treated cells (FigureS5J&K). These data are consistent with elevated actin and Src-family kinase activity reducing the rate of export of phosphorylated YAP1. Together, these data reveal that perturbation of actin and Src-family kinases alters YAP1 regulation within the nucleus; specifically, dissociation from chromatin and export. This contrasts with the expectation of decreased nuclear import of actin and of Src-regulated pathways modulating the interaction of YAP1 with cytoplasmic partners.

### Import/Export machinery screen identifies XPO1 as key mediator of YAP1 export

The different results above indicate that nuclear export is a key step in the regulation of YAP1. To learn more about the regulation of YAP1 entry and exit from the nucleus we performed a siRNA screen targeting the known complement of nuclear import and export machinery. The siRNA library was targeted against human genes, we therefore carried out the screen in two human cancer-associated fibroblasts. FigureS6A&B confirm that, similar to murine CAFs, nuclear YAP1 localization in human VCAF4 and VCAF8 depends upon actomyosin function. To identify regulators of both import and export, we sought to identify conditions in which the levels of YAP1 in the nucleus and cytoplasm were roughly equivalent. FigureS6C shows that increasing the confluence especially for VCAF8 led to similar YAP1 staining intensity in both nucleus and cytoplasm. VCAF4&8 were reverse transfected with human siRNA pools targeting 143 different genes. After 4 days we fixed and stained the cells for YAP1 and images were acquired using a Cellomics Arrayscan (Figure6A). Figure6B shows that siRNA targeting YAP1 and MST1/MST2 have the expected effect reducing overall YAP1 levels and increasing nuclear YAP1, respectively. The images were then assessed in a double-blinded manner for either altered nuclear or cytoplasmic distribution of YAP1: the overlapping cell bodies of the fibroblasts at confluence precluded accurate automated cell segmentation and determination of nuclear/cytoplasmic ratios. Several siRNA consistently perturbed YAP1 localization; TableS2 summarizes this analysis. 14 genes were selected for a secondary screen using three different siRNA distinct from those used in the original screen. XPO1, RANBP3, ZPF36, and HRB were found to consistently, and with multiple siRNA, increase the nuclear localization of YAP1 (Figure6B&C). In contrast, THOC3 decreased the nuclear localization of YAP1. A tertiary screen in the murine NF1 and CAF1 fibroblasts highlighted the central role of XPO1 in YAP1 nuclear export (Figure6D). Quantification revealed an increased YAP1 nuclear localization with approximately 4 times (NF1) and 2 times (CAF1) more YAP1 in the nucleus compared to the cytoplasm (Figure 6E). The level of XPO1 knock-down was confirmed both at the mRNA and protein level in NF1 and CAF1 (FigureS6D&E). An off-target effect due to the use of a siRNA pool was excluded by checking YAP1 localization using single oligos (FigureS6F). We next explored if XPO1 depletion would affect TEAD-driven transcription. Figure6F shows that depletion of XPO1 leads to a small, but significant, increase in TEAD-dependent transcription in NF1. A less pronounced change was observed in CAF1, most likely because YAP1 is already predominantly nuclear in control CAFs (Figure6F). These data demonstrate that XPO1 is a key mediator of YAP1 export in both human and murine cells. Further, nuclear accumulation of YAP1 in normal fibroblasts is sufficient to modestly increase TEAD-dependent transcription – albeit not as dramatically as mutation of LATS1/2 phosphorylation sites (Figure1G).

**Figure 6:**
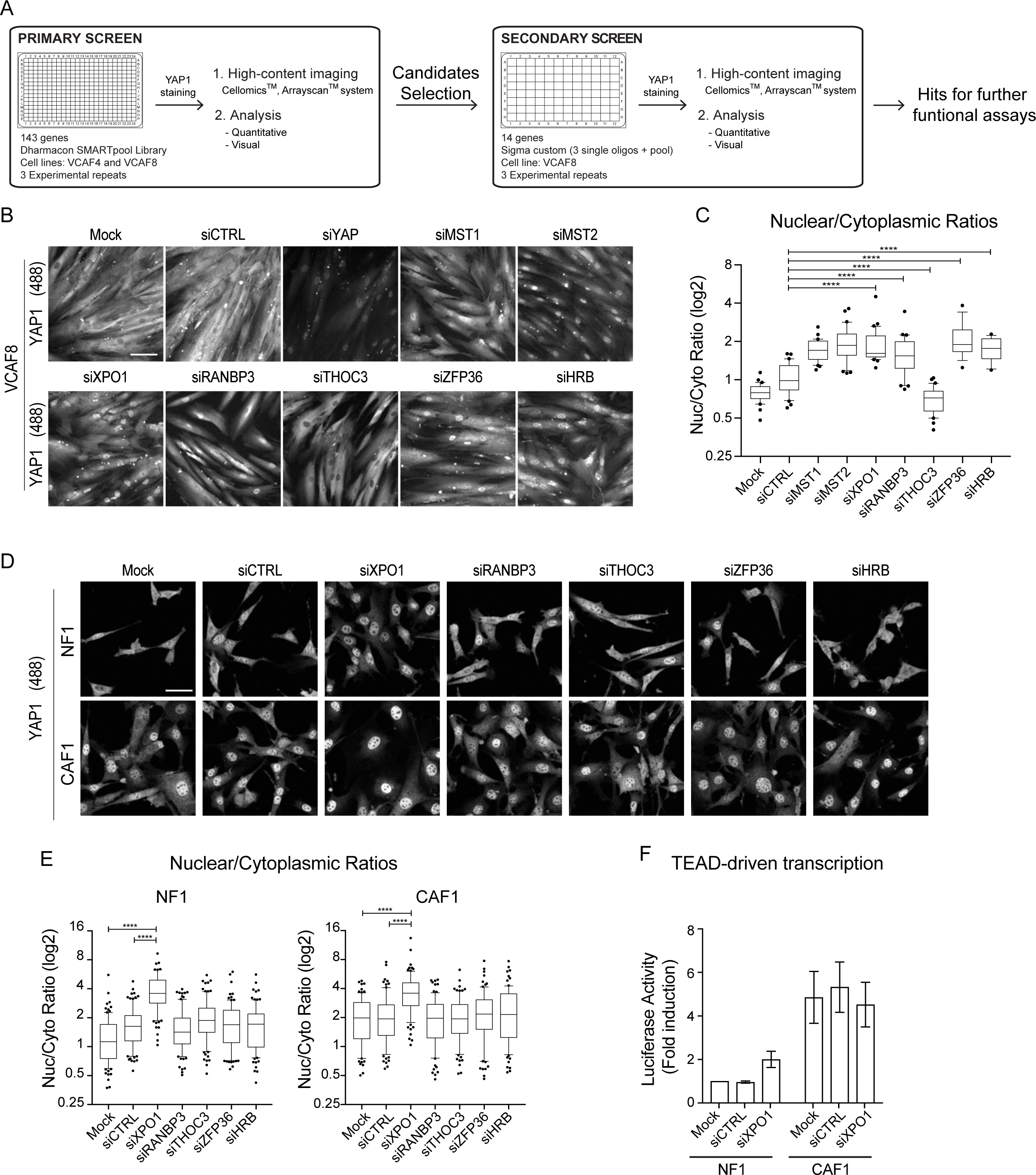
Import/Export machinery screen identifies XPO1 as key mediator of YAP1 export. (**A**) Schematic representation of the screen experimental pipeline. (**B**) Representative images of the secondary screen showing endogenous YAP1 staining in human VCAF8 upon siRNA knockdown of controls targets and several hits. Scale bar, 100μm (**C**) Box-plot (10&90) of nuclear-to-cytoplasmic ratio (log2 scale) corresponding to one experimental repeat of the secondary screen. n > 15 cells. (**D**) Representative images showing the effect of specified hits on endogenous YAP1 localization in murine cells, NF1 and CAF1. Scale bar, 50μm (**E**) Box-plot (10&90) of nuclear-to-cytoplasmic ratio (log2 scale) corresponding to three experimental repeats in NF1 and CAF1. n>89 cells. (**F**) Luciferase assay of NF1 and CAF1 upon knock-down of specified siRNA. Bars represent mean ± s.e.m. of 3 independent experiments. Mann-Whitney U-test, n.s., non significant, ** p≤0.01, **** p≤0.0001. See also Figure S6.

### Nuclear YAP1 is regulated by actomyosin

We next wished to determine the relationship between XPO1-mediated export and cytoskeletal regulation of YAP1. Figure7A&B shows that blebbistatin treatment is unable to lead to the cytoplasmic translocation of YAP1 if XPO1 is depleted. This, together with data in Figure5, argues that cytoskeletal regulation is reducing YAP1 nuclear export. Finally, we asked whether inhibiting export prevented cytoskeletal regulation of YAP1 transcriptional function. Strikingly, TEAD-dependent reporter assays indicated that blebbistatin still reduced the transcription competence of YAP1, even though it remained in the nucleus (Figure7C). These data clearly suggest that two mechanisms exist by which the cytoskeleton and Src-family kinases influence YAP1 activity. First, they reduce the export of LATS1/2-phosphorylated YAP1 thereby controlling its sub-cellular localization. Second, as shown in Figure5, cytoskeletal integrity influences the tyrosine phosphorylation of YAP1 required for its maximal transcriptional competence. Thus, divergent mechanisms downstream of Src-family kinases couple cytoskeletal integrity to YAP1 function.

**Figure 7:**
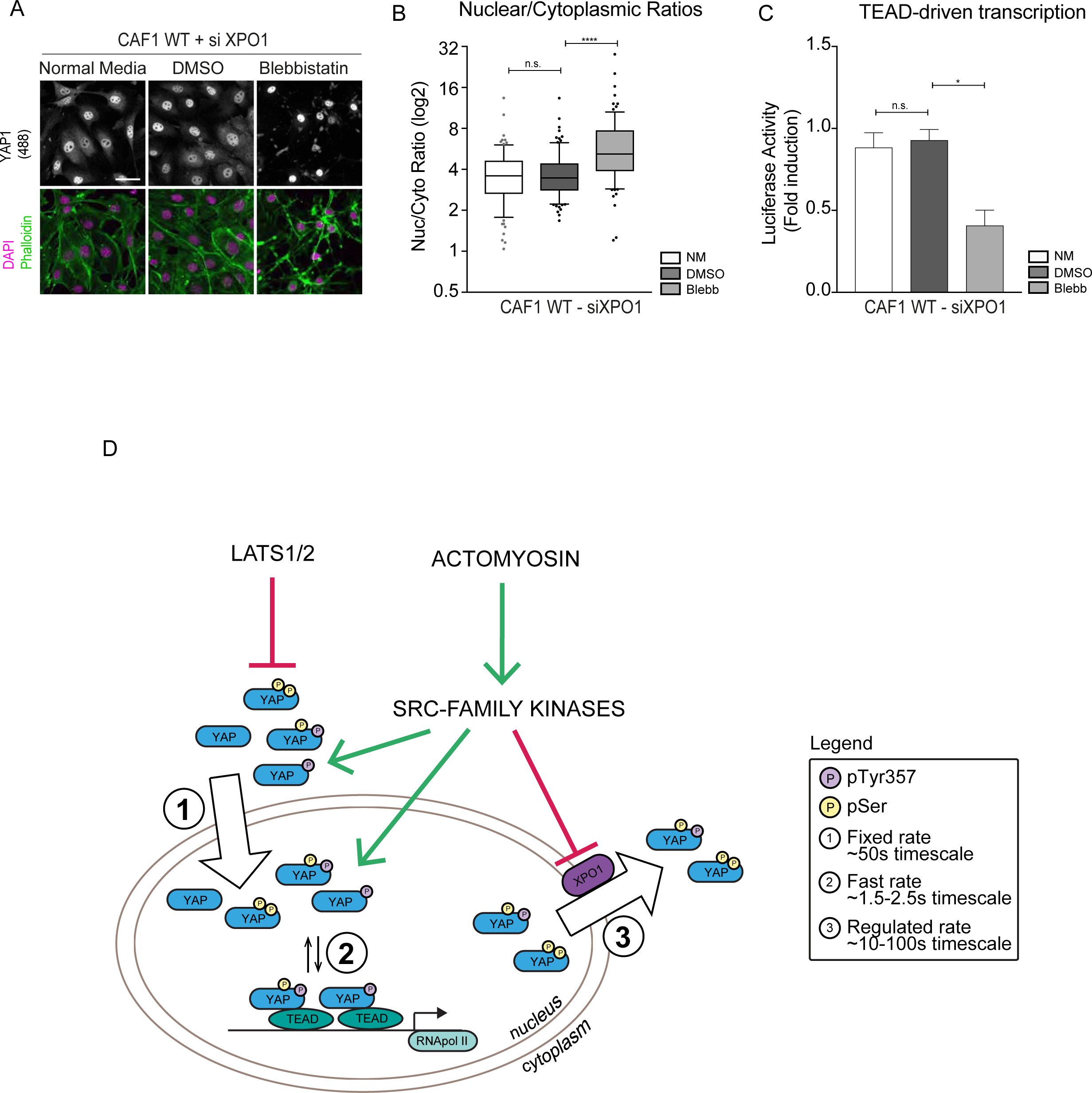
Nuclear YAP1 is regulated by actomyosin. (**A**) Representative images of endogenous YAP1 localization upon XPO1 depletion followed by blebbistatin treatment. (**B**) Box-plot (10&90) of nuclear-to-cytoplasmic ratio (log2 scale) corresponding to (A). n<80 cells from three experimental repeats. Mann-Whitney U-test, n.s., non significant, **** p≤0.0001. (**C**) Luciferase activity of CAF1 upon XPO1 depletion in normal condition and treated with DMSO or Blebbistatin. Bars represent mean ± s.e.m. of 3 independent experiments. Unpaired, t-test, n.s., non significant, * p≤0.05. (**D**) Schematic of YAP1 dynamics between the nucleus and the cytoplasm: serine phosphorylations are represented in yellow and tyrosine phosphorylations in lilac.

## Discussion

The transcriptional regulator YAP1 is critically important for the control of growth of epithelial tissues and organ size (Piccolo et al., 2014; Zhao et al., 2011; Zhao et al., 2007). YAP1 also plays a key role in fibroblast activation in pathological contexts (Calvo et al., 2013). Notably the tumor-promoting functions of cancer-associated fibroblasts depend upon YAP1, but not TAZ. Further, mesenchymal knock-out of YAP1 results in embryonic lethality around E11.5 (Ege *et al.*, in preparation). The regulation of YAP1 can be divided into a ‘canonical’ pathway involving negative LATS1/2-mediated serine phosphorylation events and a more recently described pathway linked to cytoskeletal integrity and Src-family kinase function (Low et al., 2014). Our previous work demonstrated that actomyosin function acts upstream of Src-family kinases to regulate YAP1 in CAFs (Calvo et al., 2013). However, the interplay between these pathways and how they regulate the sub-cellular dynamics of YAP1 is not understood. In this work, we have used photobleaching of fluorescently tagged YAP1 combined with molecular perturbations and mathematical modeling to tackle this issue. The use of ordinary and partial differential equation based methods present many benefits over the traditional ‘t1/2’ (half-time) and immobile fraction analysis historically used for FRAP analysis. t1/2 metrics completely overlook the pssibility of multiple processes occurring simultaneously and the estimation of the immobile fraction relies on judging asymptotic points with noisy data in systems that may not even exhibit such behavior. Our FRAP analysis enables numerical diffusion and reaction rates to potentially be determined. Furthermore, the implementation of PDE analysis of FLIP data avoids the subjective selection of reporting points for analysis and provides parameters estimates that robustly describe spatial variability.

Our quantitative analysis reveals several surprises (Figure7D). First, YAP1 is highly dynamic with molecules shuttling in and out of the nucleus in a timescale of 50-100 seconds. Second, the interaction with TEAD is a very short-lived with dissociation rates ∼0.5 s^-1^ for wild-type YAP1 and only ∼0.2s^-1^ for the strong gain of function 5SA mutant. This contrasts with the DNA binding of TEAD, which has a dissociation rate two orders of magnitude slower. Together, these data indicate that a timeframe of a second is required for YAP1 to trigger the molecular events that promote RNA polymerase II dependent transcription. The relative transience of this interaction may relate to the engagement of TEAD-occupied enhancers with the more proximal core machinery of promoters (Galli et al., 2015; Zanconato et al., 2015). The third striking observation is the dominance of nuclear export as a point of YAP1 regulation. Intriguingly, we find that serine phosphorylation is required for nuclear export. EYFP-YAP1_5SA has a greatly reduced export rate and its localization remains nuclear even when the cytoskeleton or Src-family kinase functions are perturbed. This realization represents an important shift in the view that LATS1/2-phosphorylated YAP1 is stably sequestered in the cytoplasm. Instead, the increased cytoplasmic localization of LATS1/2- phosphorylated YAP1 is the result of increased nuclear export. In agreement with this view, pS127 YAP1 is observed in the nucleus (Figure4F).

However, serine phosphorylation is not the only factor influencing YAP1 localization. Targeting actin or Src-family kinase function in CAFs increases the export rate to similar levels to those in normal fibroblasts without changing LATS1/2-mediated YAP1 phosphorylation. Thus, instead of cytoskeletal integrity releasing YAP1 from a cytoplasmic anchor, we propose that nuclear import is constitutive. This could occur via a YAP1/MASK complex with the ankyrin repeats of the latter protein generating a RaDAR nuclear entry signal (Lu et al., 2014; Sansores-Garcia et al., 2013). This step is not regulated by the cytoskeleton, instead, the cytoskeleton modulates XPO1-mediated nuclear export. The exact mechanism by which this is achieved remains to be investigated; we can exclude Y357 phosphorylation, but it remains possible that other tyrosine phosphorylation sites in YAP1 might modulate the activity of the putative nuclear export sequences (NES) in YAP1. Bioinformatic analyses suggest several possible NES in YAP1, but with very low confidence score and not consistently between the two tools used (LocNES and NesNES1.1). An alternative possibility is that emerin phosphorylation may be involved. This protein links the inner nuclear membrane to the nuclear lamina and is subject to Src-mediated phosphorylation following nuclear stress (Tifft et al., 2009). Further, it has been implicated in regulating the sub-cellular distribution of two other transcriptional regulators, β-catenin and MKL1/MRTF (Guilluy et al., 2014; Markiewicz et al., 2006).

Comprehensive functional analysis of the known nuclear import and export machinery identified XPO1 as key for the export of YAP1. This is consistent with previous work using leptomycin B, which can inhibit XPO1 (Nishi et al., 1994). Interestingly, we found no functional conservation in murine cells of numerous other hits in the human system (even though they were validated with multiple siRNA). This may reflect a multitude of minor secondary export mechanisms that are highly variable between different fibroblast lineages or species. The final striking observation is that control of sub-cellular distribution is not sufficient to strongly activate YAP1. We find that tyrosine phosphorylation of Y357 represents an independent mechanism of YAP1 regulation. It does not affect sub-cellular localization, but greatly reduces the transcriptional competence of YAP1. It is also intriguing to speculate where in the cell Src-family kinase mediated phosphorylation of this residue occurs. The simple expectation would be that this occurs, like the majority of Src-mediated phosphorylation, in the cytoplasm (Parsons and Parsons, 2004). However, the ability of the cytoskeleton to influence YAP1 transcriptional activity even when export is blocked suggests that phosphorylation may also occur in the nucleus. There are several precedents for nuclear Src activity, including the aforementioned phosphorylation of emerin and regulation of estrogen receptor (Castoria et al., 2012). Taken together these analyses lead to a reevaluation of YAP1 regulation (Figure7D). We find no evidence that it is stably sequestered in the cytoplasm. Instead, it cycles frequently in and out of the nucleus and is subject to extensive control of its rate of export. This requires ‘canonical’ LATS1/2 phosphorylation, but the rate is negatively tuned by actomyosin and Src-family kinase activity.

**Figure S1:**
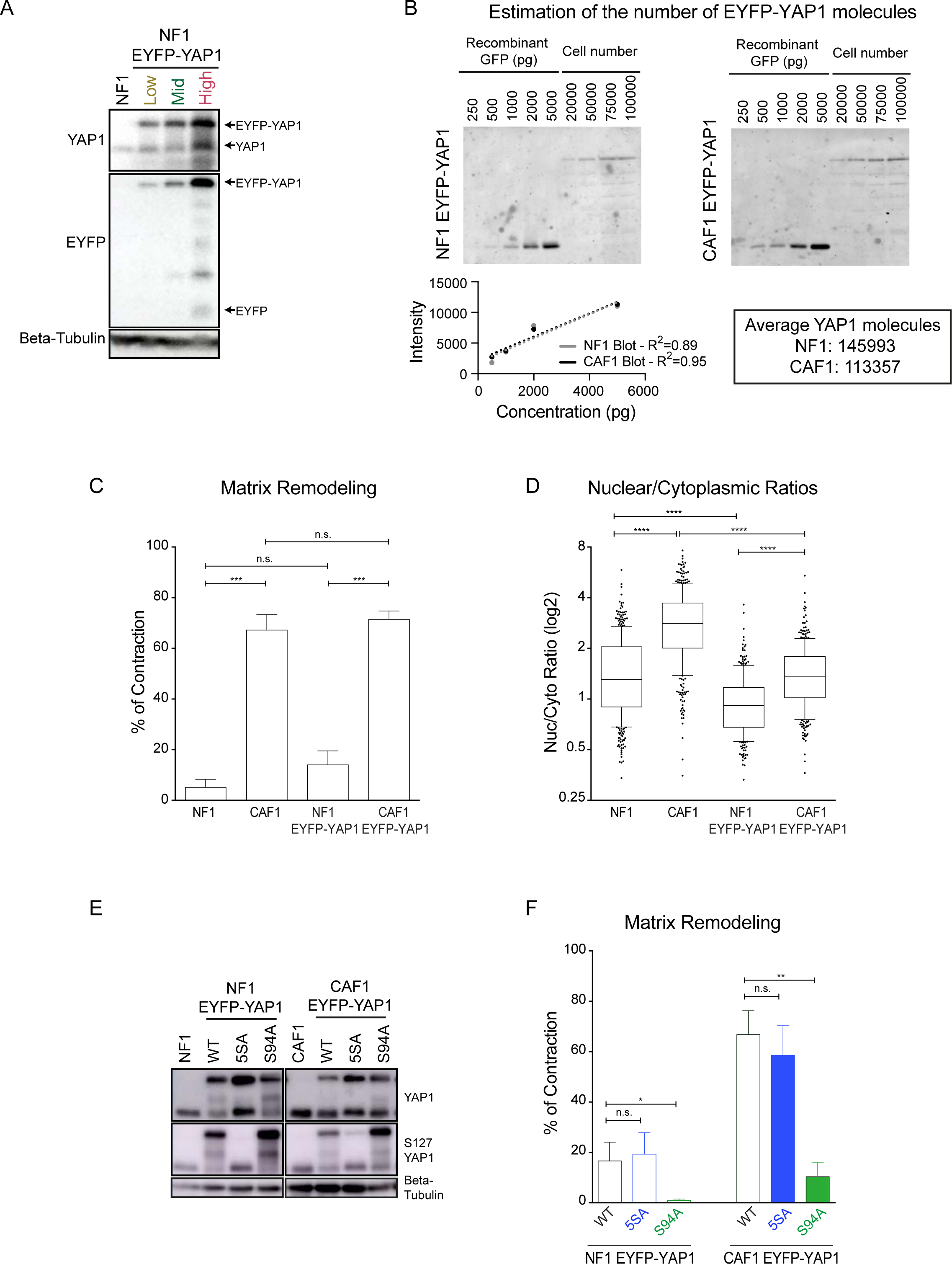
Overexpression of EYFP-YAP1 maintains fibroblasts phenotype. Related to Figure1. (**A**) Western blot showing the expressions of EYFP-YAP1 (92kDa) versus endogenous YAP1 (61kDa), in NF1 WT and in three different FACS-sorted populations (Low-Mid-High) of NF1 EYFP-YAP1. EYFP (27kDa) and Beta-Tubulin (42kDa) are also presented. (**B**) Western Blot and linear regression analyses to assess the number of EYFP-YAP1 molecules in both NF1 and CAF1, using different concentration of GFP recombinant proteins. (**C**) Contraction assay of WT and EYFP-YAP1 expressing NF1 and CAF1 in normal condition. Bars represent mean ± s.e.m. of 5 independent experiments. (**D**) Box-plot (10&90) of nuclear-to-cytoplasmic ratio (log2 scale) of endogenous YAP1 and EYFP-YAP1 in NF1 and CAF1. n>170 cells for each conditions from at least 3 independent experiments. (**E**) Western Blot showing the expressions of EYFP-YAP1 WT, 5SA, S94A (92kDa) versus endogenous YAP1 (61kDa), in NF1 and CAF1. Phosphorylation of Serine 127 and Tubulin (42kDa) are also presented. (**F**) Contraction assay of EYFP-YAP1, EYFP-YAP1_5SA and EYFP-YAP1_S94A expressing NF1 and CAF1. Bars represent mean ± s.e.m. of 5 independent experiments. Mann-Whitney U-test, n.s., non significant, * p≤0.05,** p≤0.01, *** p≤0.001.

**FigureS2:**
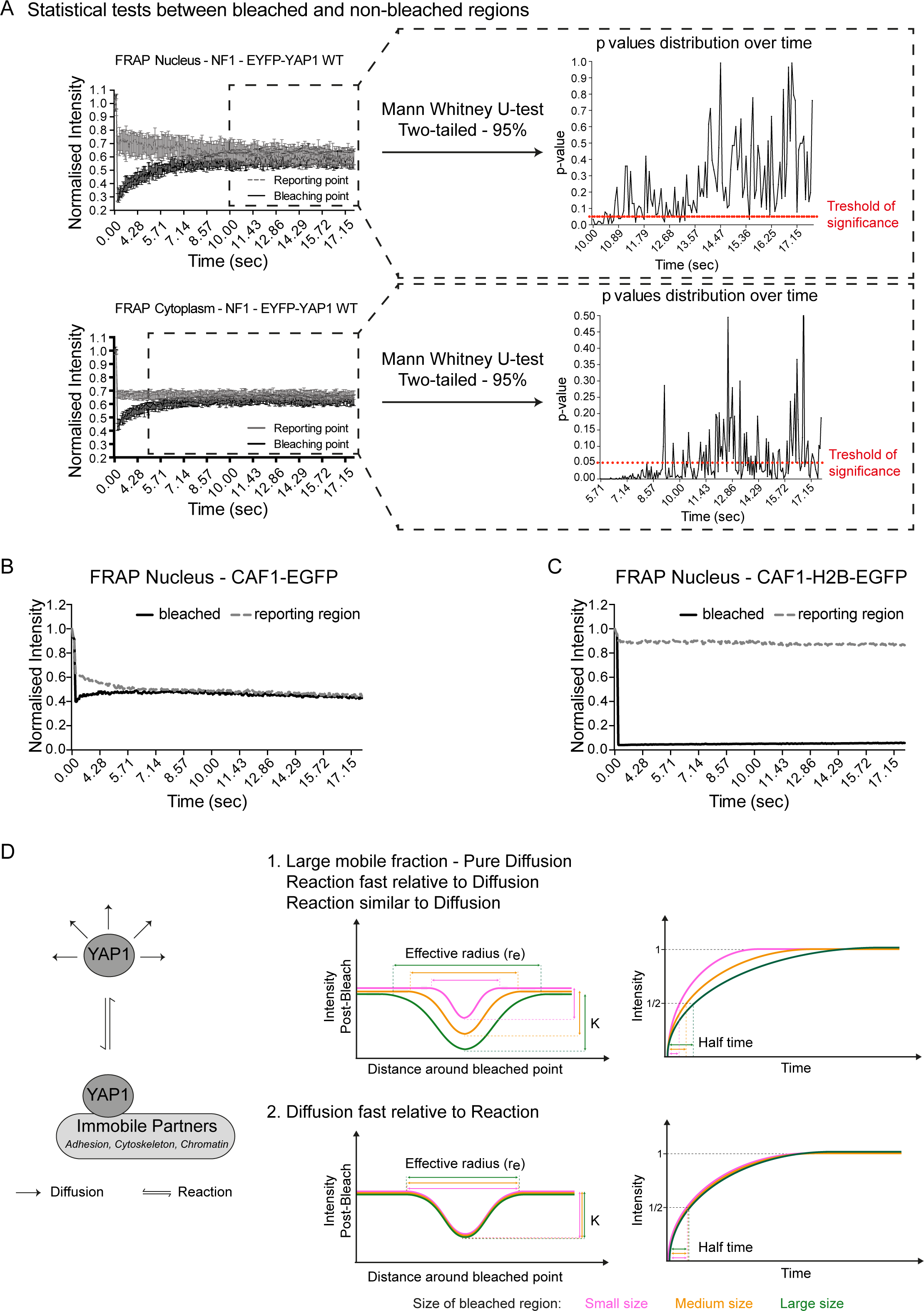
FRAP experiments to identify diffusion and nuclear dissociation rates. Related to Figure2. (**A**) Graphs showing the evolution of the p values assessing the statistical differences between the distribution of bleached and reporting EYFP-YAP1 intensities upon nuclear and cytoplasmic FRAP in NF1. FRAP graphs (left) represent median with 95%CI. (**B**) Graph showing the median of EGFP intensities from bleached (plain line) and reporting (dotted line) regions in 5 representative cells upon nuclear FRAP in CAF1. (**C**) Graph showing the median of H2B-EGFP intensities from bleached (plain line) and reporting (dotted line) regions in 5 representative cells upon nuclear FRAP in CAF1. (**D**) Schematic showing effective radius (r_e_), bleach-depth (K) and half-time plots in two contexts: 1. When diffusion is quantifiable or 2. When diffusion is too fast to be estimated. For more details, refer to Mathematical Methods. Mann-Whitney U-test, n.s., non significant.

**FigureS3:**
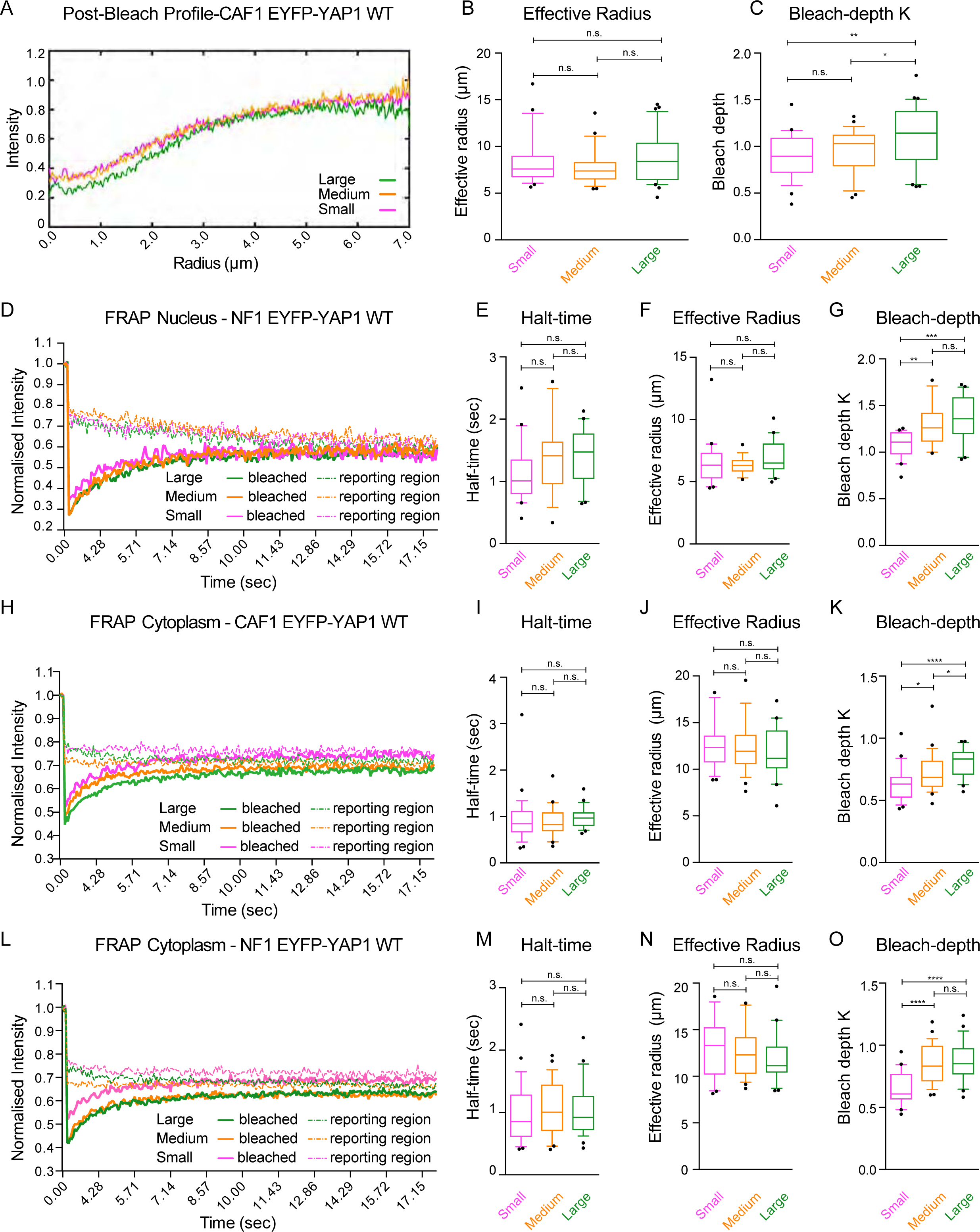
Diffusion is fast and does not take part in the recoveries observed upon FRAP. Related to Figure2. (**A**) Post-bleach profiles corresponding to (Figure2B) recoveries of intensities in CAF1 EYFP-YAP1 nuclear FRAP. (**B**) Box-plot (10&90) showing effective radius corresponding to (Figure2B) recoveries of intensities. (**C**) Box-plot (10&90) showing bleach-depth K corresponding to (Figure2B) recoveries of intensities. (**D**) Graph showing the median intensities of EYFP-YAP1 for three different sized bleached (plain line) and reporting (dotted line) regions upon nuclear FRAP in NF1 EYFP-YAP1. n= 20 cells for each of the sizes from 3 biological replicates. (**E-G**) Box-plot (10&90) showing half-time, effective radius and bleach-depth corresponding to (D) recoveries of intensities. (**H**) Equivalent graph to (D) upon cytoplasmic FRAP in CAF1, n= 30 cells for each small and medium sizes, 25 cells for large size from 3 biological replicates. (**I-K**) Box-plot (10&90) showing half-time, effective radius and bleach-depth corresponding to (H) recoveries of intensities. (**L**) Equivalent graph to (D) upon cytoplasmic FRAP in NF1 EYFP-YAP1. n= 30 cells for each of the sizes and from 3 biological replicates. (**M-O**) Box-plot (10&90) showing half-time, effective radius and bleach-depth corresponding to (L) recoveries of intensities. Mann-Whitney U-test, n.s., non significant, * p≤0.05, ** p≤0.01, *** p≤0.001, **** p≤0.0001.

**FigureS4:**
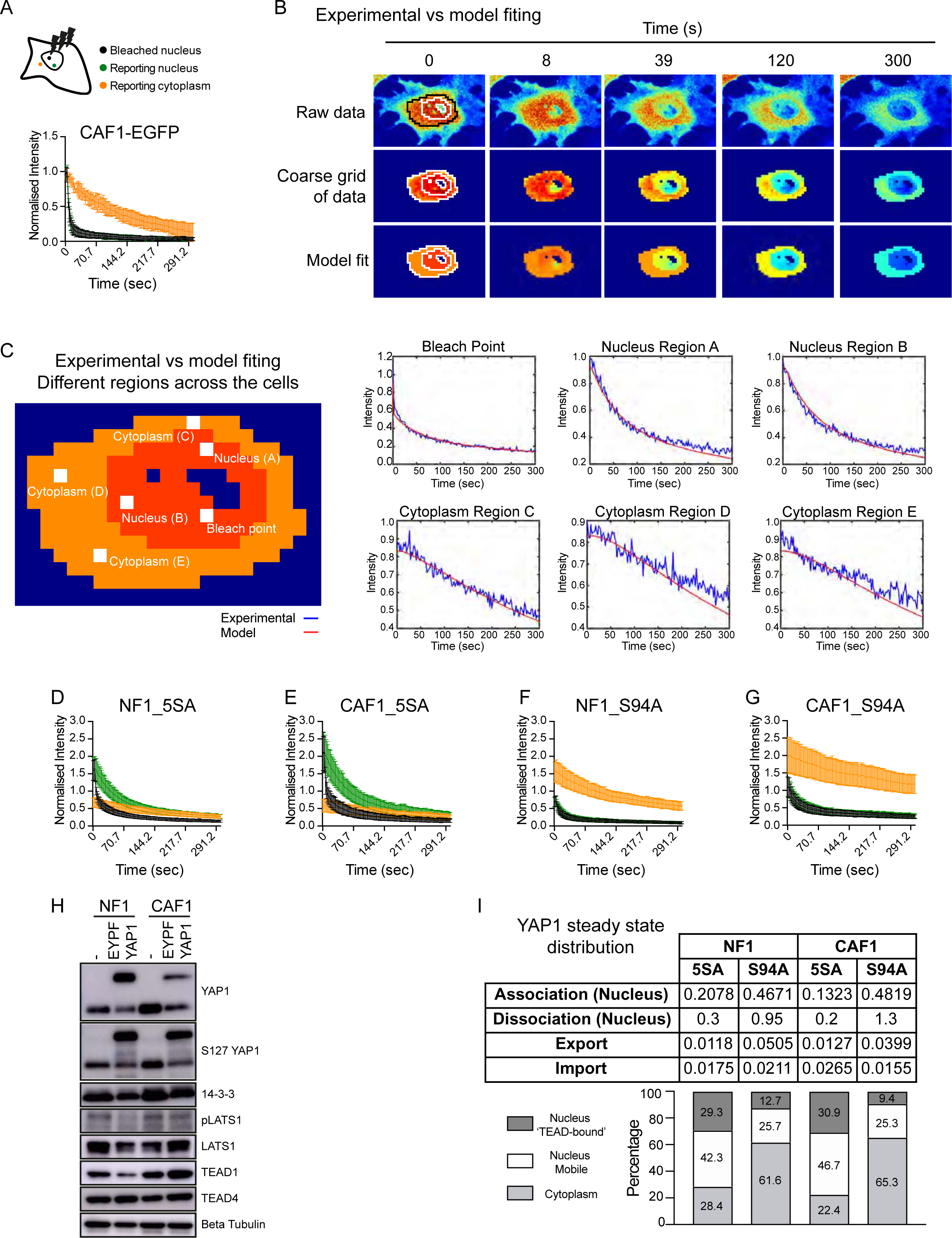
Mathematical modelind allows spatial analyses during FLIP experiments. Related to Figure4. (**A**) Graph showing the evolution of EGFP intensities from bleached (black), nuclear reporting (green) and cytoplasmic reporting (orange) upon nuclear FLIP in 5 representative CAF1 cells. Graph represents mean with 95%CI. (**B**) Comparison of experimental data to FLIP model fitting for a single CAF1 cell undergoing FLIP in the nucleus. The raw experimental data, the coarse-gridded discretization of the experimental data and FLIP PDE model fit to the coarse grid discretization are compared for various time-points (0s, 8s, 39s, 120s and 300s). The manually determined boundaries of the cytoplasm, nucleus and nucleoli are illustrated at 0s (see Mathematical Methods). (**C**) Plots of intensity versus time at the bleached point and various labeled points in the nucleus and cytoplasm for the same cell as in (B). Together B and C illustrate both good spatial and temporal fits to the experimental data. (**D**) Equivalent graph to (A) showing the evolution of EYFP-YAP1 5SA intensities upon nuclear FLIP in CAF1, n=26 cells from 3 biological replicates. (**E**) Equivalent graph to (A) showing the evolution of EYFP-YAP1_5SA intensities upon nuclear FLIP in NF1, n=30 cells from 3 biological replicates. (**F**) Equivalent graph to (A) showing the evolution of EYFP-YAP1_S94A intensities upon nuclear FLIP in NF1, n=27 cells from 3 biological replicates. (**G**) Equivalent graph to (A) showing the evolution of EYFP-YAP1_S94A intensities upon nuclear FLIP in CAF1, n=25 cells from 3 biological replicates. (**H**) Western blot of both WT and EYFP-YAP1 expressing NF1 and CAF1. YAP1 (61 and 92kDa), S127 YAP1 (61 and 92kDa), 14-3-3 (27kDa), pLATS1 (140kDa), LATS1 (140kDa), TEAD1 (50kDa), TEAD4 (48kDa), and Beta-Tubulin (42kDa) are represented. (**I**) Mathematical estimation of YAP1 steady-state distribution in NF1 and CAF1 expressing 5SA and S94A mutants.

**FigureS5:**
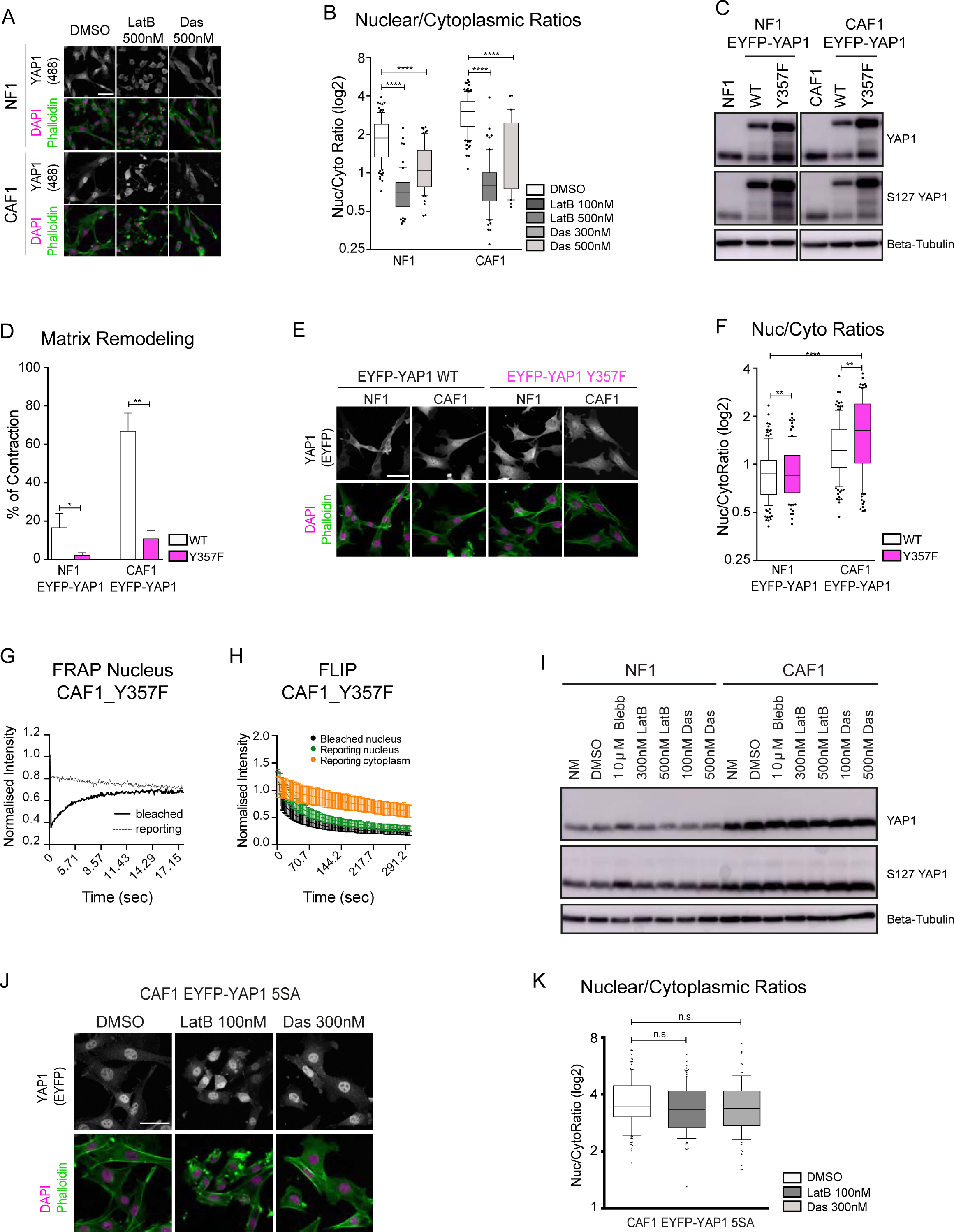
Effect of treatment with latrunculin B and dasatinib and of Y357F mutation. Related to Figure5. (**A**) Representative images of endogenous YAP1 localization in NF1 and CAF1 treated with DMSO or latrunculin B and dasatinib. Scale bar, 50μm. (**B**) Box-plot (10&90) of nuclear-to-cytoplasmic ratio (log2 scale) of endogenous YAP1 in NF1 and CAF1 treated with DMSO or latrunculin B and dasatinib, n>30 cells from at least 2 independent experiments. (**C**) Western Blot showing the expressions of EYFP-YAP1 and EYFP-YAP1_Y357F (92kDa) versus endogenous YAP1 (61kDa), in NF1 and CAF1. Phosphorylation of Serine 127 and Tubulin (42kDa) are also presented. (**D**) Contraction assay of NF1 and CAF1 expressing EYFP-YAP1 vs EYFP-YAP1_Y357F. Bars represent mean ± s.e.m. of 5 independent experiments. (**E**) Representative images of EYFP-YAP1_Y357F localization in NF1 and CAF1. Scale bar, 50μm. (**F**) Box-plot (10&90) of nuclear-to-cytoplasmic ratio (log2 scale) of EYFP-YAP1 vs EYFP-YAP1_Y357F localization in NF1 and CAF1, n>100 cells from at least 3 independent experiments. (**G**) Graph showing the median intensities of EYFP-YAP1_Y357F from bleached (plain line) and reporting (dotted line) regions upon nuclear FRAP in CAF1, n= 30 cells from 3 biological replicates. (**H**) Graph showing the intensities of EYFP-YAP1_Y357F from bleached (black), nuclear reporting (green) and cytoplasmic reporting (orange) regions upon nuclear FLIP in CAF1, n= 28 cells from 3 biological replicates. Graph represents mean with 95%CI. (**I**) Western blot showing the effects of latrunculin B and dasatinib treatment on S127 YAP1 in NF1 and CAF1 WT. (**J**) Representative images of EYFP-YAP1_5SA in CAF1 treated with DMSO or latrunculin B and dasatinib. Scale bar, 50μm. (**K**) Box-plot (10&90) of nuclear-to-cytoplasmic ratio (log2 scale) of endogenous EYFP-YAP1_5SA in CAF1 treated with DMSO or latrunculin B and dasatinib. n>90 cells from 3 independent experimental repeats. Mann-Whitney U-test, n.s., non significant. Mann-Whitney U-test, n.s., non significant, * p≤0.05, ** p≤0.01, **** p≤0.0001.

**FigureS6:**
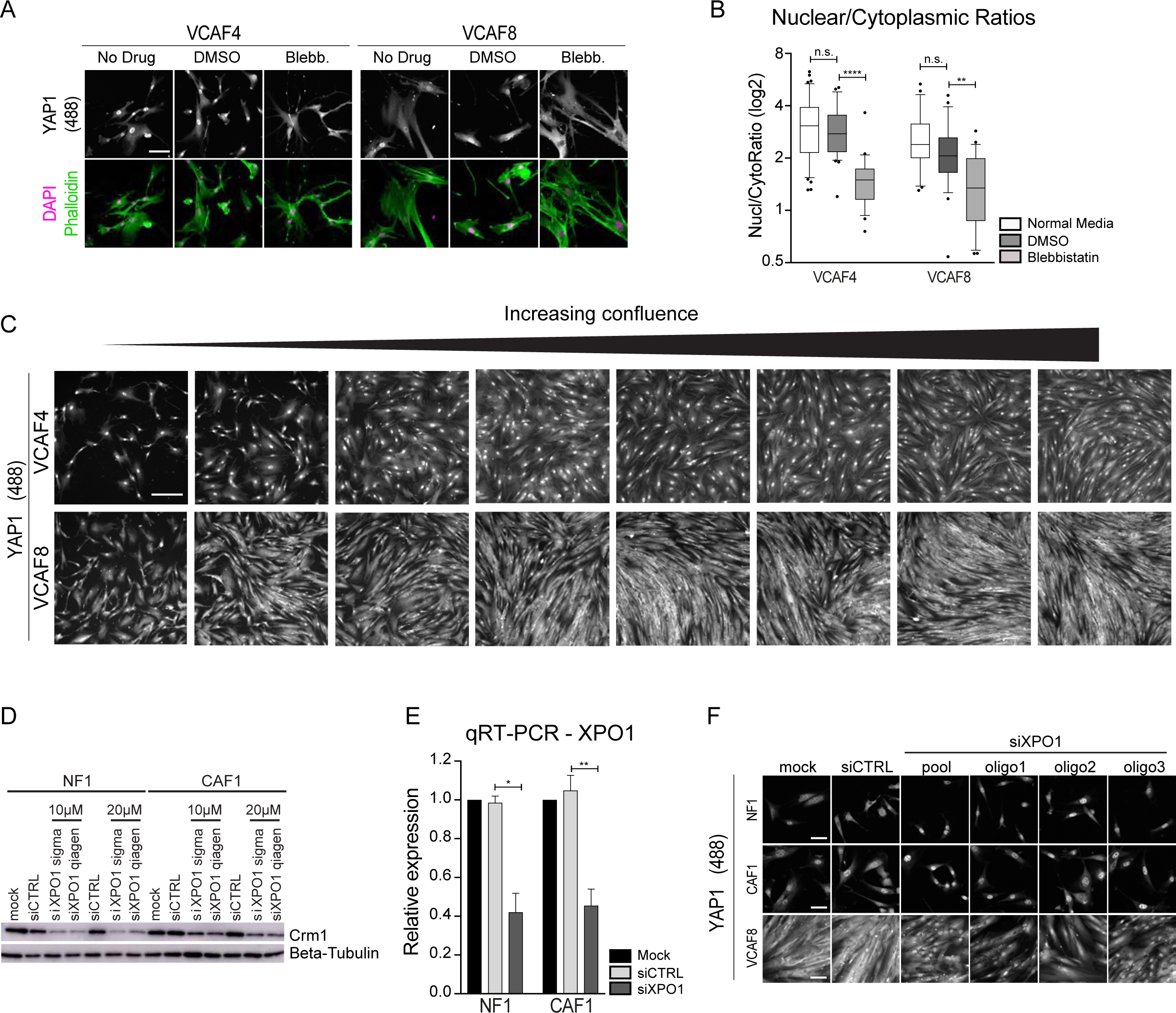
Validation of conditions and reagents for the siRNA screen. Related to Figure6. (**A**) Representative images showing endogenous YAP1 localization in human VCAF4 and VCAF8 cell line treated with blebbistatin. Scale bar, 100μm. (**B**) Box-plot (10&90) of nuclear-to-cytoplasmic ratio (log2 scale) corresponding to data in n > 21 cells from one experimental repeat. Mann-Whitney U-test, n.s., non significant, ** p<0.01, **** p<0.0001. (**C**) Representative images of endogenous YAP1 staining in human VCAF4 and VCAF8 cell lines with increasing cell confluence. Scale bar, 200μm. (**D**) Western blot showing depletion of Crm1 (123kDa) in NF1 and CAF1 upon XPO1 depletion with different concentration and reagent. Beta-Tubulin (42kDa) are also presented. (**E**) qRT-PCR showing the level XPO1 depletion in NF1 and CAF1 upon depletion with 20μM with Sigma reagent. Bars represent mean ± s.e.m. of 3 independent experiments. Unpaired, t-test, n.s., non significant, * p≤0.05. (**F**) Representative images showing endogenous YAP1 localization in NF1, CAF1 and VCAF8 upon XPO1 pool deconvolution to single oligos. Scale bar, 50μm.

**TableS1: Related to Figure1:** Table compiling output of mathematical models fitting experimental data for nuclear FRAP of NF1 and CAF1 expressing EYFP-YAP1.

**TableS2: Related to Figure2:** Result of the primary siRNA screen in humand CAFs.

**TableS3: Methods:** List of Antibodies, siRNA and primers for QRT-PCR and cloning.

**Movies:** Stars indicate when photo-bleaching is performed at the end of the frame.

Movie1: Nuclear FRAP of CAF1 expressing H2B-GFP and EGFP. Scale, 4 μm.

Movie2: Nuclear FRAP of NF1 and CAF1 expressing EYFP-YAP1. Scale, 4 μm.

Movie3: Cytoplasmic FRAP in NF1 and CAF1 expressing EYFP-YAP1. Scale, 4 μm.

Movie4: Nuclear FRAP of NF1 and CAF1 expressing either EYFP-YAP1_5SA or EYFP-YAP1_S94A. Scale, 4 μm.

Movie5: Nuclear FRAP of NF1 and CAF1 expressing TEAD1-mCherry. Scale, 4 μm.

Movie6: FLIP of NF1 and CAF1 expressing EYFP-YAP1. Scale, 10 μm.

Movie7: FLIP of NF1 and CAF1 expressing either EYFP-YAP1_5SA or EYFP-YAP1_S94A. Scale, 10 μm.

Movie8: FRAP of CAF1 expressing EYFP-YAP1_Y357F or EYFP-YAP1 treated with latrunculin B and dasatinib. Scale, 4 μm.

Movie9: FLIP of CAF1 expressing EYFP-YAP1_Y357F or EYFP-YAP1 treated with latrunculin B and dasatinib. Scale, 10 μm.

## Methods

### Cell lines

Normal mammary glands fibroblasts (NF1) and mammary carcinoma fibroblasts (CAF1) are described in (Calvo et al., 2013). Briefly, fibroblasts were isolated from transgenic FVB/n MMTV-PyMT mice and immortalised with HPV-E6 retrovirus. Human vulval (VCAF) fibroblasts were isolated from patient tissue samples collected from patients at Bart’s and the London Hospital under ethical approval 10/H0304/14 and immortalised by pBABE-Hygro-HTERT retroviral transfection. All EYFP-YAP1 constructs (described in Plasmids section) were introduced into the cells using the Lentivirus system. Upon transfection, populations expressing EYFP signal were selected by fluorescence-activated cell sorting (FACS). All cells were cultured in DMEM (Invitrogen), 10% FCS (PAA Labs), 1% ITS (insulin-transferrin-selenium; #41400-045; Invitrogen).

### Plasmids

Both Lenti-EF-EYFP-YAP1 and Lenti-EF-mCherry-TEAD1 were generated using LentiLox 3.7 vector (pLL3.7 – gift from Way lab) where CMV promoter was replaced by EF1 alpha promoter. EYFP-YAP1 fusion was cloned from pDEST EYFP-YAP1 plasmid, gift from Nic Tapon, Francis Crick Institute. The EYFP fluorophore was well-suited to our photobleaching experiments because it is not as photostable as some more ‘optimized’ derivatives of GFP, but is sufficiently bright to be easily imaged (Shaner et al., 2005). YAP1 corresponds to the human YAP1-2γ isoform, which is 504 amino acids long (Gaffney et al., 2012; Sudol, 2013). The different mutants were generated by site-directed mutagenesis. The five serine mutated for the 5SA mutant correspond to serines 61, 109, 127, 164 and 397 (=381) for the isoform used in this study. The tyrosine ‘357’ corresponds to the residue 407 in the isoform used in this study. mCherry-TEAD1 was generated fusing TEAD1 from pRK5-Myc-TEAD1 plasmid (Addgene, ♯33109) to mCherry. EGFP and H2B-EGFP constructs were generated previously in the lab and are both cloned under CMV promoter. The two plasmids used for the luciferase experiments, pGL3-49 and pGL3-5xMCAT-49, a gift from by Nicolas Tapon.

### Immunofluorescence

All immunofluorescence experiments were performed on cells seeded on glass in a 35 mm glass bottom MatTek dish (P35-1.5-14-C, MatTek Co., Ashland, MA, USA). Cells were fixed in 4% paraformaldehyde, washed with PBS and permeabilised by incubation in 0.2 % Triton X100, PBS for 3-5 minutes at room temperature. Samples were subsequently blocked for 1 hour with 5%BSA, PBS before incubation with primary antibody for YAP1 (Santa Cruz, sc101199, 1:200) in 3%BSA, PBS overnight at 4°C. Primary antibody was washed off in 3 washes of 15 minutes with 0.05 % Tween-20, PBS. Fluorescent secondary antibody (Life Technologies) was diluted 1:500 in 3%BSA, PBS and incubated with the samples for one hour, then washed off with 3 washes of 0.05 % Tween-20, PBS. Samples requiring staining for F-actin or the nucleus were also stained with 633-phalloidin (SIGMA) and DAPI (SIGMA) at 1:500 dilution. After 3 washes of 15 min in PBS, secondary antibody in blocking solution was added. After 3 washes of 15 min in PBS, cells were imaged using an inverted Zeiss LSM710/780/880 inverted confocal laser scanning microscopes equipped with an argon laser (Zeiss, Germany).

### Photobleaching experiments

Cells were plated at low confluence and cultured overnight in 35 mm glass bottom MatTek dishes (P35-1.5-14-C, MatTek Co., Ashland, MA, USA) in DMEM media with 10 % FCS and 1% ITS. One hour prior imaging, the media was changed to Leibovitz L-15 media (CO2 independent – Invitrogen) −1% serum. The cells were subsequently bleached and imaged with a Zeiss LSM880 inverted confocal laser scanning microscope equipped with an argon laser (Zeiss, Germany) using 514nm laser and a 63X objective (Zeiss, alpha-Plan Apochromat 63x/1.46 NA oil korr TIRF).

For FRAP (Fluorescent Recovery After Photobleaching) experiments, three sizes of circular ROIs were used: small, medium and large. Small, medium and large corresponds to 11×11 pixels (3.1 μm^2^), 14×14 pixels (4.6 μm^2^) and 17×17 pixels (6.9 μm^2^) in the nucleus, respectively and 14×14 pixels (4.6 μm^2^), 17×17 pixels (6.9 μm^2^) and 20×20 pixels (9.4 μm^2^) in the cytoplasm, respectively. All images were 8-bit and 128×128 pixels. Before photobleaching, 3 measurements of fluorescence were taken. The ROI was then photobleached for 2.9 seconds using maximum laser power. A series of images were then taken every 60 milliseconds for up to 18 seconds, enough to observe complete recovery for most conditions. In order to have complete recovery of intensity EYFP-YAP1 5SA samples were imaged up to 48 seconds. FRAP of TEAD1-mCherry were imaged for 100 seconds. Graphs for FRAP represent only the 17×17 pixels size, except for diffusion analyses where all three sizes are represented.

For FLIP (Fluorescent Loss In Photobleaching) experiments, a single square ROI of 8×8 pixels (4.4 μm^2^) was used. All images were 12-bit and 128×128 pixels. Before photobleaching, 3 measurements of fluorescence were taken. The ROI was then photobleached between every frame for 2 seconds using maximum laser power. A series of 150 images were taken every 2 seconds for up to 5 minutes.

### Image analysis

For quantification of subcellular localization of EYFP-YAP1, the nuclear-to-cytoplasmic ratio was calculated. Using MetaMorph (Molecular Devices), the integrated intensities in three square regions of interest (ROI) of the same size, 8×8 pixels (11 μm^2^) were measured: one region in the nucleus, one in the cytoplasm and one outside any cells to assess the background intensity. The two ROIs in the cells were positioned at equal distance from the nuclear boundary. To normalise, the background intensity was subtracted from the two other intensities. Subsequently, the nuclear intensity was divided by the cytoplasmic intensity in order to find the nuclear-to-cytoplasmic ratio.

For quantification of FRAP (Fluorescent Recovery After Photobleaching) experiments, the integrated intensities were followed using MetaMorph (Molecular Devices). Three circular ROIs of the same size as the bleached ROI were followed: bleached ROI, one reporting ROI in the bleached nucleus and one ROI outside any cells to measure the background intensity. In parallel, the intensity of the whole field was measured to follow loss of intensity due to multiple and continual acquisition. To normalise, first the background intensity was subtracted from all other intensities. Then, nuclear intensities were normalized to the loss of intensity of the whole field over the entire experiment. Finally, they were normalized to their intensity in the first frame, prior to photobleaching. It is the recovery of this normalised intensity that was plotted and analyzed quantitatively (see Mathematical Modeling methods).

For quantification of FLIP (Fluorescent Loss In Photobleaching) experiments, MetaMorph (Molecular Devices) was used to follow the integrated intensities of six different square ROIs of the same size as the bleached ROI: the bleached ROI, two reporting ROIs (one in the same compartment as the bleached point and one in the other compartment), two controls ROIs (one in the nucleus and one in the cytoplasm of one control cell) and one ROI outside any cells to measure the background intensity. The two nuclear ROIs and the cytoplasmic ROI of the bleached cells were positioned to be at approximately equal distances form one another. To normalise, first the background intensity was subtracted from the five others intensities measured in the cells. Then, the intensities measured in the cell of interest (three ROIs) were normalised by dividing by the average intensity of the two control ROIs. These data were plotted in the graphs. Our mathematical model – used to extract import rates, export rates and association rates – was based on the intensities of the whole cells rather than 6 individual ROIs (see Mathematical Modeling Methods). For all the image analyses using ROIs, the nucleoli regions, where YAP1 appeared to be excluded, were avoided.

### Luciferase assay

Luciferase assays were performed with the dual luciferase assay kit (Promega). Cells were lysed using passive lysis buffer. Lysates were placed into a white 96-well plate (Perkin Elmer) to assess Luciferase and Renilla activities using Envision Multilabel plate reader (Perkin Elmer). To normalise, the measurements of firefly luciferase activities were normalised to the renillla luciferase activities of the same sample.

### ECM remodeling assay

To assess force-mediated matrix remodeling, 50 × 10^3^ fibroblasts were embedded in 100 μL of Collagen I:Matrigel (#354249: #354234; BD Biosciences) and seeded on a 35 mm glass bottom MatTek dish (P35-1.5-14-C, MatTek Co., Ashland, MA, USA). Once the gel was set, cells were maintained in DMEM + 10% FCS + 1% ITS, unless otherwise stated. Gel contraction was monitored daily by taking photographs of the gels. The gel contraction value refers to the contraction observed after 2 days. To obtain the gel contraction value, the relative diameter of the well and the gel were measured using ImageJ software, and the percentage of contraction was calculated using the formula 100 × (well diameter – gel diameter) / well diameter.

### Transfections, inhibitors and siRNA

Primary and secondary screens were done on human fibroblasts using reverse siRNA transfection with Lipofectamine RNAiMAX (#13778030, Thermo Fisher), where VCAF4 and VCAF8 cells were added directly to wells containing siRNA (384/96well plate for primary/secondary screen respectively) using manufacturers instructions. For primary screen, 1000 cells were seeded per well, while for secondary screen 3500-4000 cells were seeded. Cells were fixed 4 days after transfection and stained with YAP/TAZ antibody and DAPI. For siRNA transfection in mouse fibroblasts (NF1 and CAF1), reverse transfection method was used for siRNA transfection with Lipofectamine RNAiMAX (50pmol siRNA per well, 6 well plate) using 60000-75000 cells per transfection. For DNA transfection, cells were subject to transfection with Lipofectamine LTX and Plus reagent (# 15338100, Thermo Fisher) the following day (2.5μg DNA per well, 6 well plate). When specified, drug treatment was started 6hrs after DNA transfection, lasting 48hrs. siRNA was purchased from Dharmacon or Sigma, and sequences are listed in Supplementary Table2). The following drugs were used: Blebbistatin – 10μm (#203391; Calbiochem /Merck), Dasatinib – 300nM or 500nM (LC Labs), LatrunculinB −100nM or 500nM (LC Labs).

### Western Blot

All protein lysates were obtained, processed and ran following standard procedures. Antibody description and working dilutions used can be found in Supplementay Table2.

### Statistical analysis

Statistical analyses were performed using Prism software (GraphPad Software). p values were obtained using Mann-Whitney unpaired t–test or paired-t-test with Welch’s Correction, with significance set at p < 0.05. Graphs show symbols describing p values: *, p < 0.05; **, p < 0.01; ***, p < 0.001, ****, p<0.0001.

### Mathematical model

Please refer to Mathematical Methods.

## Author Contributions

Conceptualization, N.E. and E.S.; Methodology, N.E., R.P.J., and E.S.; Formal analysis, N.E. and R.P.J.; Investigation, N.E., A.M.D., M.J., and M.H.; Resources, M.H.; Writing – Original Draft, N.E., R.P.J., and E.S.; Supervision, E.S.

## Acknowledgments

All authors were funded by Cancer Research UK and The Francis Crick Institute. We thank Dr Nic Tapon for providing us with plasmids used in this study. We also thank Reuben O’Dea for discussion and feedback on the mathematical aspect of this study. We finally thank lab members for help and advice throughout this work.

## 1 Mathematical Methods

### 1.1 FRAP Data Analysis

Estimation of the reactive and diffusive processes taking place during FRAP included analysis of both the postbleach intensity profile of the cell frame following the bleach process and the dynamic recovery of intensity in the Region Of Interest (ROI). Mathematical derivation leading to this methodology can be found in Kang *et al.* (2009, 2010) and it explicitly accounts for rapid diffusion of the protein of interest over the timescale used for photobleaching. We briefly explain the main points of data fitting here. Methods for image acquisition and intensity normalization in specified ROIs are explained in the main methods.

#### 1.1.1 Postbleach Profile Analysis

The prebleach (the frame captured prior to the bleach process taking place) and postbleach profiles (the frame immediately captured after completion of the bleach process) were re-centred around the mid-point of the nominal bleach region (Supplementary Figures Ia-b with the red circle in (Ia) illustrating the bleach region). An image mask of the nucleus minus the nucleoli regions was created by manually tracing the relevant boundaries using MATLAB’s ‘roipoly’ command with nucleoli most obvious from the pre-bleach profile (Supplementary Figure Ic). This image mask enabled us to consider intensity changes only in the nucleus. Nucleoli were excluded because the dense chromatin packing excludes YAP and alters protein diffusion. Both the postbleach profile and image mask were transformed from Cartesian to polar coordinates using MATLAB’s ‘cart2pol’ function (Supplementary Figures Id – e). Data-points were then interpolated over the polar angle, *θ*, to account for the lower density of points near the centre of the nominal bleach region (Supplementary Figure If). The median postbleach profile was then calculated as the median intensity of all data-points within the image mask for increasing radius from the nominal bleach point (blue curve in Supplementary Fig Ig). It was confirmed that an exponential of a Gaussian (red curve)

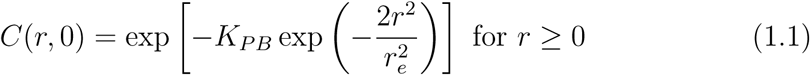

fit the median postbleach profiles well where *r* is the radial distance from the origin, *r_e_* is the effective radius (measure of distance along x-axis in (Ig)) and K_PB_ is the bleach-depth (measure of drop in intensity on y-axis in (Ig)). By minimizing the sum of squares due to error, the parameters *r_e_* and *K_PB_* for which the exponential of a Gaussian that best fits the data could be determined.

#### 1.1.2 Recovery Curve Analysis

Three possible model fits to the recovery curve, *S(t*), were then compared ( Sprague *et al.* (2004)). These were i) a diffusion model arising from either the mobile fraction being large (pure diffusion) or the on/off binding rates being fast relative to diffusion (effective diffusion); ii) a reaction-diffusion model that makes no assumptions on relative rates of binding and diffusion and iii) a reaction model where diffusion is assumed to be rapid and the recovery is determined by the slow reaction from immobile to mobile state. Each of the models thus incorporate some combination of rates of transfer from bound to unbound states (dissociation), rates of association from unbound to bound states (association) and rates of diffusion in bound and unbound states. These parameters are given respectively by *k_on_*, *k_off_* and D for association, dissociation and diffusion.

##### Pure Diffusion and Effective Diffusion Models

In addition to being derived from the postbleach profile (1.1), the bleach depth can alternatively be calculated via the recovery curve intensity. Utilizing the point of completion of the bleach process, *S(0*) in the recovery curve, the corresponding bleach depth, *K_RC_*,solves the equation

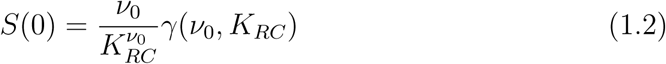

where 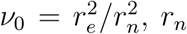 is the nominal bleach radius i.e. the radius of the bleach region and 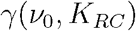 is the incomplete gamma function. The two parameters estimating bleach depth, *K_PB_* and *K_RC_*, may not be exactly equal. For model fitting to the recovery curve, Kang *et al.* (2009) recommend using the value derived from the recovery curve itself, for fitting consistency.

The diffusion function, *Q_D_ (t*), fitted to the recovery curve *S(t*) is then given by

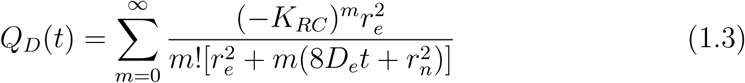

where *D_e_*, the only free parameter in this function is selected to minimize the weighted residuals (1.6) between this function *Q_D_(t*), and the recovery curve data, *S(t*). The diffusion coefficient is thus obtained explicitly via knowledge of the effective radius, *r_e_*, in the postbleach profile.

##### Reaction-diffusion Model

For the reaction-diffusion model, in addition to the parameters required for the diffusion only model, two additional parameters are determined from the recovery curve: *f_0_*, the normalized initial postbleach intensity (a value between zero and one) and R, the mobile fraction of recovery, calculated as

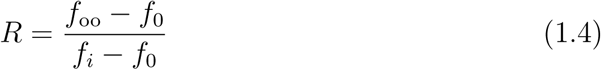

where *f_oo_* gives the mean intensity of the recovery curve data, once it has reached steady-state, and *f_i_* gives the mean intensity of the recovery curve prior to bleaching (due to normalization, this value will be equal to or close to one). The reaction-diffusion function, *Q_RD_(t*), fitted to the recovery curve is rather more complex than the others described here (see Kang *et al.* (2010) for further details). The corresponding MATLAB m-file, BDfrap.m was used to calculate these reaction-diffusion function values for varying *k_on_*, *k_off_* and *D_1_* (diffusion corresponding to the unbound state). We assumed that diffusion *D_2_* corresponding to the bound state was *0μm^2^s^-1^* (supported by the FRAP data of H2B which does not diffuse – Supplementary Figure S2C) to reduce the degrees of freedom in the model fit. The selection of the unknown parameters *k_on_*, *k_off_* and *D_1_* was then again made to minimize the weighted residuals between this function and recovery curve data.

##### Reaction Models

Reaction models were fitted as described in Fritzsche & Charras (2015) such that

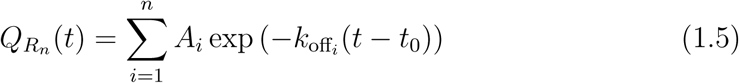

where *n =1* for the single reaction and n = 2 for the double reaction. Here, *A_i_* gives the amplitude for recovery, k_offi_ the corresponding rate of recovery and t_0_ allows the model fitting to account for noise in measurement at time zero of the recovery – the postbleach intensity.

##### Weighted Sum of Squares of Error

Without loss of generality, let the function, *Q(t)*, refer to each of the reaction-diffusion based models described. Data fitting was then carried out to minimize the time-weighted residuals between the data, *S(t*), and model *Q(t*) as described in Kang *et al.* (2009) such that data-points for earlier time contribute more to the residual, allowing greater capacity to identify faster rates. We minimize the time-weighted sum of squares of error, given by

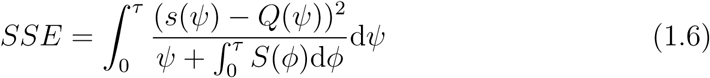

where *τ* is the final point in time of the data and the integral in the denominator is included to remove the singularity at t = 0.

##### Implementation of Model Fitting

The nonlinear regression of each proposed model, *Q(t*), to the data, *S(t*), was carried out using the MATLAB algorithm ‘nlinfit’, found in the Statistics and Machine Learning Toolbox. The approach uses the Levenberg-Marquardt nonlinear least squares algorithm to find the fit that minimizes the weighted SSE (1.6). Initial parameter guesses are required and the algorithm then searches for the global minimum of the weighted SSE using these initial guesses as starting points. Poor initial guesses can lead to the algorithm becoming slower in reaching the global minimum or, even worse, becoming stuck in a local rather than global minimum. Care should therefore be taken in the choice of these initial guesses. Fortunately, the Levenberg-Marquardt approach is quite robust to poor initial parameter guesses, although it is then slower in reaching the minimum.

The diffusion parameter was initially guessed at *19μm^2^s^-1^*. We estimated that EYPF-YAP1 has a volume roughly four times greater than a single GFP. We thus interpolated the rate of diffusion of three and five GFPs through the nucleus as estimated by Baum *et al.* (2014). In order to provide good initial guesses for association and dissociation (and related amplitudes of recovery) for the models incorporating reaction dynamics the exponential function

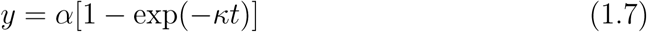

was fitted to each recovery curve whereby the fits for *α* and *κ* could be used as guesses for amplitude and association/dissociation for each curve. The function (1.7) is also nonlinear and so to derive *α* and *κ* we used the nlinfit algorithm and again needed initial guesses. For a small subsample of cells, a grid was constructed for the two parameters *α* and *κ* and the standard SSE calculated at each point on the grid. This identified the region of parameter space where the global minimum occurred as being *α* ≈ 0.3 and *κ* ≈ 0.5. For the fit of (1.7) to each curve using the nlinfit algorithm we could then use 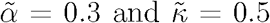 as initial parameter guesses. The output values for *α* and *κ* estimated from the nlinfit algorithm were then used as initial guesses for various parameters in the models *Q_RD_(t*) and *Q_R_n__(t*).

For reaction-diffusion, *Q_RD_ (t*), the initial guesses were given by 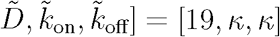. In the case of the single reaction, *Q_R_1__(t*), the initial guesses were given by 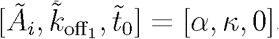. For the double reaction, the initial rates are estimated similarly to Fritzsche & Charras (2015). The fast reaction is assumed to contribute the first 30% of overall recovery and the slow reaction contributes the final 30% of recovery. We estimate where these regions occur based on the single reaction fit, *Q_R_1__(t*), (due to it being a smooth curve as opposed to the original noisy data).

To determine the model that most likely describes the behavior of the system we make use of the effective radius and bleach depth derived from the postbleach profile alongside a number of statistics that describe the model fitting to recovery curve. The postbleach statistics implied that diffusion was likely to be fast and that we were in a regime where the observed recovery was dominated by slower reactions. From the model fitting to the recovery curves, we first compared the four model fits using the Akaike Information Criterion (AIC), a statistic that rewards goodness of fit but penalizes models based on the number of parameters required for that fit. From the AIC values for the four models, the Akaike weights (normalized relative likelihoods of each model) are calculated. These Akaike weights can be interpreted as the probability that a certain model is most likely, given the data. Importantly, the AIC analysis does not say that a specific model is correct, it says only how likely the model is to be true in comparison to the other models. For each model in which the Akaike weights suggested that diffusion or reaction-diffusion were most likely we then considered the fitted parameters for that model. Model fits whereby the fitted diffusion parameter was fitted at over *60μm^2^s^-1^* were ruled out as erroneous as they approached or exceeded the measured diffusion rate in cells of a single GFP which is approximately a quarter of the size of EYPF-YAP1 (see Baum *et al.* (2014)). Similarly, reaction-diffusion models in which the binding on and/or off rates were greater than *25s^-1^* were again determined to be erroneous. Such rates would be incredibly rapid and would not be able to be accurately estimated due to our frame-rate. A small number of fits were recorded as diffusion or reaction-diffusion. In cases where Akaike weights or implausible parameter values ruled out diffusion and reaction-diffusion models the single and double reaction models were interpreted. Firstly, the Akaike weights for both the single and double reactions were compared. Secondly, an F-test comparing nested models was carried out on the two reaction models, with F-statistic given by

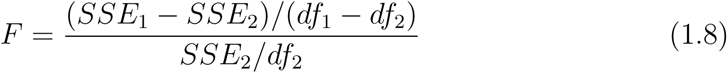

with *df*1 — *df*2 and *df*2 degrees of freedom. Here *SSE*_1_ and *SSE*_2_ are the time-weighted sum of squares of error (1.6) of the single and double reaction models respectively and *df*_1_ and *df*_2_ are the degrees of freedom of the single and double reaction models respectively. Significant p-values (< 0.05) alongside low Akaike weights for the single reaction implied a difference between the two models suggesting that a double reaction may be more likely than a single reaction. Nonsignificant p-values (*p* > 0.05) alongside Akaike weights for the single reaction of a similar magnitude to (or larger than) the double reaction suggested either that there was not enough evidence to assume a double reaction or strong evidence to assume a single reaction. In either case, a single reaction was assumed. Finally, if the F-test and Akaike weights implied that a double reaction was most likely, the fitted parameters were considered with double reactions with on/off rates greater than *25s^-1^* again being ruled out as erroneous. A double reaction was deemed most likely only if these rates fell below this range, otherwise a single reaction was again preferred.

### 1.2 FLIP Model Fitting

To model the FLIP data we developed a compartmentalized Partial Differential Equation (PDE) model that incorporated the bleach region, the rest of the nucleus and the cytoplasm (Supplementary Figure Ila). The PDE contains one compartment corresponding to the nucleus and another corresponding to the cytoplasm. The two compartments are linked via flux boundary conditions.

#### 1.2.1 FLIP PDE Description

Within the nucleus we assume a single binding reaction

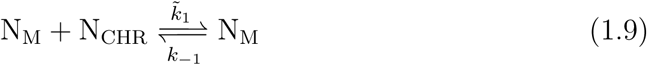

where N_M_ represents unbound mobile proteins in the nucleus, N_CHR_ the chromatin binding partners such as TEAD and N_I_ the bound immobile state. We set *N_M_(x, t), N_I_(x, t*) and *N_CHR_(x, t*) to be the respective concentrations of N_M_, N_I_ and N_chr_ at time t and location x=(*x,y).* Setting 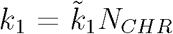, we arrive at the general PDE reaction-diffusion model within the nucleus, given by

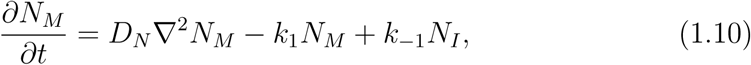

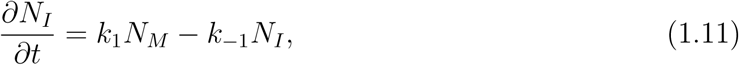

reflecting our assumption that molecules in the bound immobile state, *N_I_*, do not diffuse and remain stationary. Here, 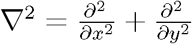. Bleaching occurs over a set region within the nucleus, which we describe with the reactions

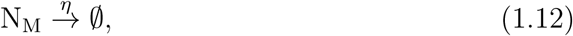

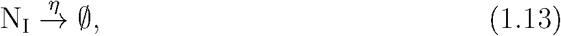

where the empty set, 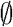, corresponds to molecules that have been bleached and effectively left our system and *η* gives the rate of bleaching. Thus, in the nucleus, we have

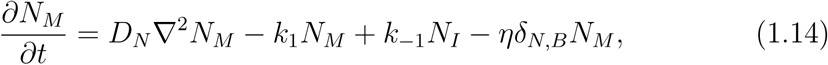

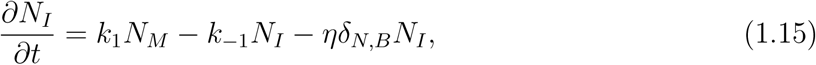

where the delta function is defined

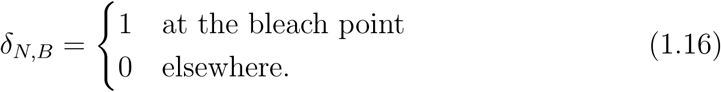

Regions of nucleoli drastically affect the mobility of the molecule within the nucleus. They are densely packed making it difficult for molecules to diffuse in or out of them. No clear YAP1 signal is observed in these regions. Hence, in our system we treat nucleoli as impenetrable by molecules and therefore impose zero-flux boundary conditions where the nucleus is in contact with these impenetrable islands. That is

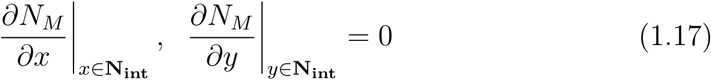

where N_int_ is the boundary between the nucleus and islands of nucleoli.

In the cytoplasmic compartment, we assume that the entire molecular population is mobile (observe the FRAP recovery times in Supplementary Figure S3) such that we obtain the single diffusive equation

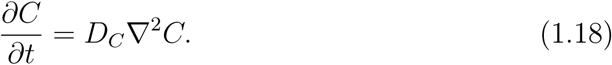

From here, we assume that the molecules diffuse at the same rate in both the nucleus and the cytoplasm such that *D = D_N_ = D_C_*. At the exterior boundary of the cytoplasm (the boundary of the cell) we impose zero-flux boundary conditions such that there is no concentration gradient at the boundary i.e.

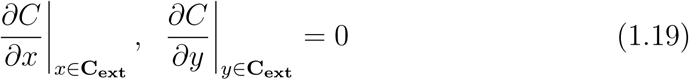

where C_ext_ is the cytoplasmic exterior.

Linking these two compartments is import and export. We can express this import/export between the two compartments via the reaction

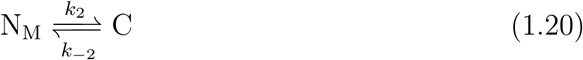

where it is assumed that only the mobile fraction in the nucleus can be exported and any molecules imported from the cytoplasm are mobile in the nucleus. These two compartments are linked via flux boundary conditions (Supplementary Figure IIa) on the exterior of the nucleus (in contact with the cytoplasm) and the interior of the cytoplasm (in contact with the nucleus). In the case of the nucleus compartment, the outward flux is then proportional to k_2_N_ext_ where N_ext_ is the nucleus exterior and the inward flux is proportional to k_-2_C_int_ where C_int_ is the internal cytoplasmic boundary. For the case of the cytoplasmic compartment, the outward flux is then proportional to k_-2_C_int_ and the inward flux to k_2_N_ext_.

#### 1.2.2 Numerical PDE fitting

##### Numerical Implementation in MATLAB

We solved the compartmentalized PDE numerically in MATLAB. Finite differences were applied in the spatial domain to account for diffusion between grid-points in the same compartment and flux boundary conditions (import and export) between the nuclear compartment and cytoplasmic compartment. This reduced the PDE to a system of ODEs for each grid-point which were solved in the temporal domain using ode15s. The ODE solver ode15s, for use in stiff problems (i.e. systems that include widely varying time-scales), was selected due to the fact that the solution has a region of very sharp change in concentration where it is not stiff alongside regions of slowly changing concentration where it is stiff. The sharp change in concentration occurs most notably in and around the bleach point at the initiation of the bleach process. The slowly changing concentration occurs both for the remainder of the cell far from the bleach point, for all time, and the region in and around the bleach point for longer time (when the majority of molecules within the cell have been bleached).

##### Image Discretization

To find the parameters in which the numerical solution to the PDE best fits our data we discretize the cell into coarse grid-points with each grid-point being the same dimensions as our bleach region, a 2.1μm^2^ square. The lattice of grid-points was initialised such that one of the grid-points overlapped exactly with the bleach point (Supplementary Figure IIb). The lattice was selected to be more computationally efficient and more robust to erroneously defined flux boundaries between the nucleus and cytoplasm. Future iterations of the model fitting will attempt to fit a finer lattice, whilst controlling for the effects of increasing noise at the boundaries, in order to further refine parameter estimates.

The nuclear, nucleola and cytoplasmic boundaries are then determined manually by the user, again using MATLABs roipoly command. In the case of the cytoplasm, we excluded regions where the cell was very thin. For example, lamellipodia are much thinner than the focal section achieved by the microscope (200-300nm compared to > 1 micron) and the low fluorescence intensity in these regions reflects the low volume of cytoplasm in the focal plane, not the differential dynamics or localization of the fluorescent protein. Typically, the cytoplasmic area selected for analysis was 1-2 times the area of the nucleus.

Image masks of both the nucleus minus nucleoli and the cytoplasm are then used to determine which grid-points are cytoplasmic and which are nuclear. Grid-points are defined as nuclear or cytoplasmic if at least 50% of that grid-point is occupied by the nuclear or cytoplasmic image mask (Supplementary Figure IIc). The intensity at a given grid-point is then calculated as the mean intensity of all pixels within that grid-point. Only pixels that overlap the nuclear or cytoplasmic masks are used in this calculation. This then defines our coarsely gridded cell with which we can fit our PDE to (Supplementary Figure IId).

##### Implementation of Model Fitting

When fitting the model to the data, a number of parameters are required to be estimated: diffusion, *D*, association *k*_l_ and dissociation *k*_-l_, import *k*_-2_ and export *k*_2_, the rate of bleaching, *η* and the initial concentrations of the bound *N_Io_* and unbound *N_Mo_* states in the nucleus and the cytoplasmic concentration *C*_0_. We fix the diffusion at *D* = 19μm^2^s^-1^ for all cases. This is an interpolation of the rate of diffusion of three and five GFPs through the nucleus reported in Baum *et al.* (2014). We fix the dissociation rate, *k*_-l_, to equal the median value for each cell-type derived from our FRAP analysis.

For the initial nuclear and cytoplasmic concentrations, we do not simply use the pre-bleach levels derived from the data due to noise in acquisition. However, we can reduce the number of free parameters by fixing the initial concentrations in terms of each other. Prior to any bleaching having occurred, the concentrations of both the nuclear and cytoplasmic compartments are assumed to be homogeneously distributed. By reducing the spatial system to a simple system of Ordinary Differential Equations (ODEs) for concentration, the transfer between the two compartments described by the import/export reaction with bound/unbound states in the nucleus can naively be expressed as

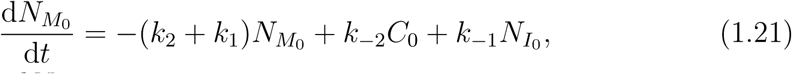

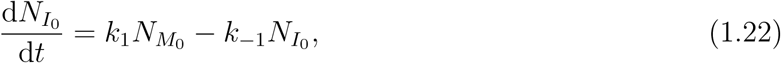

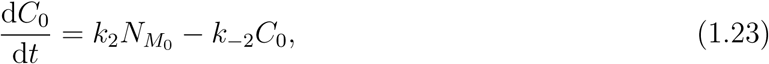

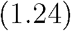

where the subscripts in N_M0_, N_I0_ and C_0_ indicate that our system is in steady-state prior to perturbations due to bleaching. Thus, at this steady-state, we observe the relations between concentrations *N_M0_* = *k_-2_C_0_/k_2_* and *N_I0_* = *k_1_N_M0_/k_-l_* and we fix concentrations in the nucleus in terms of reaction rates and cytoplasmic concentration, reducing the number of free parameters by two.

Therefore, we estimate the parameters *k_1_, k_2_, k_-2_, η* and *C_0_* by fitting our model to the data.

As with FRAP, the nonlinear regression of the compartmentalized PDE model to the data was carried out using the MATLAB algorithm nlinfit and we require initial parameter guesses for *k_l_, k_2_, k_-2_, η* and *C_0_*. For the initial cytoplasmic intensity, *C_0_*, we used the median intensity within our cytoplasmic region. For association rate, *k_l_*, our initial guess was set to be equal to the median dissociation rate acquired from FRAP for that cell type. For the final parameters, import, *k_-2_*, export, *k_2_* and decay due to bleaching, *η*, we initially set up a simpler ODE based model that incorporated just these three mechanisms and fitted to data from a single reporting point in the nucleus and another in the cytoplasm. The single reporting point in the nucleus could be either the bleach point or another location. This simple ODE model given by

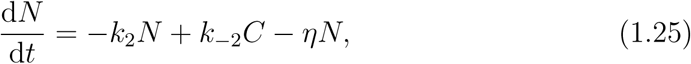

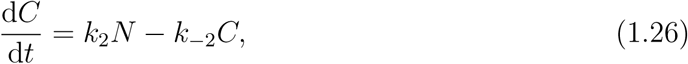

allowed us to get a naive idea of the magnitudes of import, export and decay due to bleaching via analysis of quality of fit for multiple cells. Hence we made initial guesses of export and import of around 0.002*s*^-L^ to 0.005*s*^-l^, with the nuclear/cytoplasmic ratio helping to inform on which should be initially guessed to be larger. A good initial guess for this bleaching decay was found to be 1.5*s*^-L^ via model fitting analysis of both this simpler ODE model and our full compartmentalized PDE model.

##### Weighted Sum of Squares of Error

Data fitting was carried out to minimize spatial-time-weighted residuals between the data and model. As with FRAP, data-points for earlier time contribute more to the residual. The data-points were spatially weighted such that the entire nucleus minus the bleach-point, entire cytoplasm and bleach-point were equally weighted with each other. This avoided the critical bleach-point and, to a lesser extent, the nucleus surrounding the bleach-point having a low influence on the overall model fit due to the larger size of the rest of the cell. Within the nucleus excluding the bleach point and cytoplasm each grid-point was also weighted equally with all other grid-points in that given compartment. More formally, let the signal at a given grid-point *i, j* and compartment *κ* (where *k = B* for the bleach-point, *k = N* for grid-points in the nucleus excluding the bleach point and *k δ C* for grid-points in the cytoplasm) be given by *S_i,j,k_(t*) for time-point t. We set 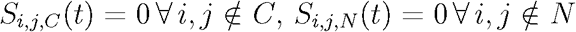 and 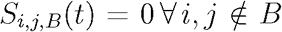. To account for both the time-weighting and equally weighted grid-points in each compartment we set

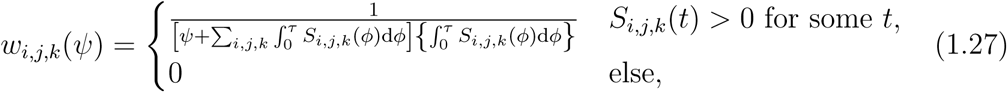

where *τ* is the final point in time of the data. The function in square brackets in the denominator is time-weighted, as in FRAP, except the integral to remove the singularity at *t = 0* is summed over all grid-points. The integral in the curly brackets in the denominator ensures each grid-point within a compartment has equal weighting on the best-fit by normalizing by the total intensity over all time of the signal at that grid-point. Finally, the grid-points are re-weighted such that the bleach-point, entire cytoplasm and entire nucleus minus the bleach point all have equal weighting with each other. Let

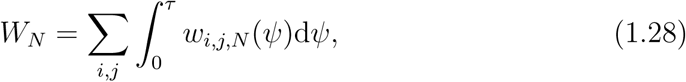

i.e. the total sum of weights of all grid-points in the nucleus for all time. Similarly, let

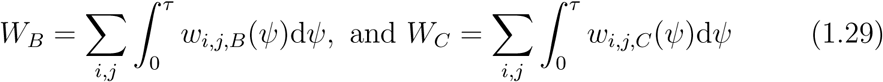

for the bleach-point and cytoplasm. Then 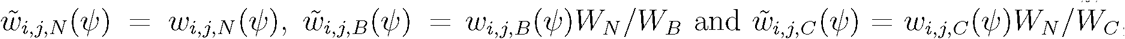, such that the nucleus minus the bleach-point, the bleach-point and the cytoplasm all confer equal weighting on the overall fit. We then seek to minimize

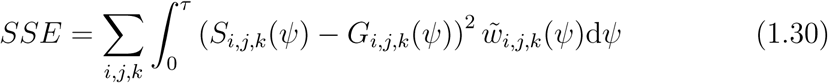

where *G_i,j,k_*() is the solution to the compartmentalized PDE at the relevant spatial grid-point and time-point.

## 2 Sensitivity Analysis

### 2.1 FRAP

#### 2.1.1 Postbleach profile

Supplementary Figure IIIa provides a heatmap of Sum of Squares due to Error (SSE) as the bleach depth, *K_PB_* and effective radius, *r_e_* are varied. The heatmap demonstrates that our choice of parameters are at the global minimum.

#### 2.1.2 Recovery Curve

Supplementary Figure IIIb provides the SSE for Equation (1.2) fitted to the recovery curve as the bleach depth, *K_RC_*, is varied. Once again, the plot demonstrates our parameter fit occurs at the global minimum. Finally, in Supplementary Figure IIIc we plot the heatmap of SSE of the fit of a single exponential reaction as the amplitude and rate of reaction are varied. Again, we found our choice of parameters occurs at the global minimum.

### 2.2 FLIP

We have carried out a sensitivity analysis over six cells, three NFWT and three CAFWT to analyze how robust our association, import and export parameter fits are to noise in our estimates for the dissociation rates (acquired from FRAP) and the rate of diffusion estimated from Baum *et al.* (2014). Here we illustrate our findings via a single NFWT and CAFWT.

Our initial analysis focusses on the sensitivity of the five free parameters, *k_l_, k_2_, k_-2_, η* and *c_0_* to the dissociation rate and diffusion rate, both fixed in the FLIP model.

The dissociation rate is acquired via FRAP analysis of each cell. However, we use a static value that does not account for intercellular variation. We begin by varying the dissociation rate and seeing how this affects the fitting of the other parameters. Predictably, the association rate is most sensitive to the value of the dissociation rate. This is due to the rate of diffusion being fixed and thus the spread of the bleach around the nucleus is reliant on the proportion of the molecule that is mobile at any point in time. For small *k*_off_ (i.e. value twice the fitted value or below) the relationship is nonlinear with the association rate decreasing faster than the association rate (Supplementary Figures IVa and IVe). This suggests that when it takes a long time for the molecule to dissociate from the bound state, the association rate needs to be even slower to allow for a greater pool of mobile protein at any point in time. For large *k*_off_ (i.e. greater than two) the relationship becomes linear, however, for fast dissociation rates, the association rate increases about three times as fast. This is due to the reverse of the above: when molecules dissociate very fast then the association rate must increase faster to allow for a greater immobile pool at any point in time. This is a consequence of the fact that the rate of diffusion is fixed. The cytoplasmic intensity, bleach decay rate and import from the cytoplasm are not sensitive to rate of dissociation. The rate of export is sensitive to rate of dissociation although much less so than the association rate. The export rate is sensitive only for low dissociation rates. For low dissociation, the export rate decreases below the fitted value. This is most likely a consequence of the proportion of mobile protein increasing (see above). As the dissociation rate increases, the mobile fraction decreases and thus the export rate must increase to fit to the data.

We now consider dissociation fixed (at 0.55*s*^-1^ for NFs and 0.40*s*^-1^ for CAFs) and observe the effects of varying diffusion on the other parameter fits (Supplementary Figures IVb and IVf). All parameters are more sensitive to very slow diffusion than other rates of diffusion. Since we have already observed that diffusion must be fast, this suggests we are in a region of parameter space where parameter fits are less sensitive to the rate of diffusion. Predictably once again, the association rate is most sensitive to the diffusion rate. For low rates of diffusion, the association rate tends to zero implying that for the model to fit well, the entire molecular population must be mobile. For faster rates of diffusion, the association rate increases, implying a smaller mobile fraction. The association rate tends to a constant for fast diffusion, demonstrating that for rapid diffusion, the association rate is not sensitive to changes in diffusivity. The effects on export rate are similar to those of association rate but export is not as sensitive to diffusion as association rate. A low diffusion rate implies the mobile fraction must be large to account for the spread within the nucleus. This in turn requires a reduction in the export rate to fit the global nuclear and cytoplasmic concentrations. The reverse is then true for fast diffusion rates. That is, export rate is dependent on the diffusion rate only via the association rate. The import, decay rate and cytoplasmic intensity are insensitive to the model fit apart from being mildly sensitive at very low rates of diffusion. This possibly reflects the model assumptions beginning to break down.

Following from this we fixed all the parameters to their fitted values and varied the association rate and export rate simultaneously. We calculated the weighted-SSE for each fit, generating a heat map of error of fit versus association rate and export rate (Supplementary Figures IVc and IVg). The results clearly demonstrate that our fitted parameters for association rate and export occur at the global minimum.

Similarly we fixed all parameters at their fitted values except for import and export. The heatmaps of weighted-SEE for import/export (Supplementary Figures IVd and IVh) demonstrate that although we are at the global minimum, it is shallower than the association rate/export rate equivalent. The heatmap demonstrates a linearity in relationship between import and export similar to the relationship between association and dissociation rates. This linearity reflects the ratio of nuclear/cytoplasmic intensity within the cell. However, the nucleus and cytoplasm have differing levels of YAP on the flux boundary between the two compartments and these levels change over time. Only the unique global minimum truly reflects these dynamically shifting values on the boundary.

Supplementary Figure IVi gives a horizontal linescan of the FLIP mathematical model output for the CAF cell shown in Figure S4H at various time-points. The results demonstrate the spatial stability of the PDE solution. The solution is relatively flat in the cytoplasm. In the nucleus the solution is flat at the boundaries with the cytoplasm and smoothly decreases towards the sink at the bleach-point. Between the nucleus and cytoplasm there is a discontinuous jump in intensity between the two compartments.

## 3 Residual Analysis

Residual Analysis has been carried out for both the FRAP and FLIP model fitting (Supplementary Figure V).

Residual analysis (observed value minus predicted value) for FRAP was carried out on the model fits for pure diffusion (Va), reaction-diffusion (Vb), single reaction only (Vc) and double reaction only (Vd) on the recovery curves of 30 NFWT (10 of each bleach radius). In each case, the red line represents the median residual of all 30 NFWT cells at each time-point. The figures demonstrate that the pure-diffusion model is inappropriate to describe the recovery as it significantly underestimates the short-time recovery and significantly overestimates the long-time recovery. However, the residuals of the reaction-diffusion, single reaction only and double reaction only models demonstrate no obvious systematic behavior, being distributed normally around zero, suggesting that each of these models describes the data well. With this in mind, the simplest system that describes the data best may be most appropriate, in this case the single reaction only model. This is in agreement with the AIC and F-test analysis (Supplementary Table1 and Section 1.1.2 above).

Residual analysis of FLIP of a single cell (Supplementary Figure Ve) demonstrates no obvious functional link between the residuals and spatial region of a cell, suggesting that the model can correctly describe the appropriate spatial aspects of the system. We then considered 15 CAF WT cells and extracted the residuals versus time for the bleach-point (Supplementary Figure Vf), remainder of the nucleus (Vg) and the cytoplasm (Vh). The results suggest that the model slightly overestimates the rate of decay due to bleaching (Vf) and as a consequence, overestimates the long-time behavior at the bleach-point. The residuals at the remainder of the nucleus and cytoplasm are generally distributed around zero. The more sparse regions of residuals in Supplementary Figures Vg and Vh could illustrate erroneous boundary determination between the nucleus and cytoplasm such that some grid-points defined as nuclear are cytoplasmic and vice-versa. This effect would become more pronouncedif there is significant cell movement. Again, for small time, the residuals do hint at the model overestimating the rate at which YAP leaves the system due to bleaching. This could be a consequence, for example, of an unaccounted for additional reaction that reduced the overall motility of YAP in the nucleus. Overall, the residual analysis suggests that the FLIP model is good but does hint at aspects of the model that could be investigated for improvement in later iterations, perhaps to identify further reactions.

## 4 Mathematical Methods Supplementary Figure Legends

### Mathematical Methods Supplementary Figure I: FRAP postbleach profile processing and analysis

Walkthrough of analysis of the postbleach profile of a single CAFWT cell undergoing FRAP. (a) Pre-bleach profile (frame prior to bleach process) re-centred such that the bleach-point (red circle) is in the centre of the image. Nucleoli are observable as regions of low intensity within the nucleus. (b) Post-bleach profile (first frame captured upon completion of bleach process) re-centred around bleach-point (red circle in (a)). (c) Image mask (re-centred around the bleach-point) outlining the manually determined boundaries of the nucleus and nucleoli. (d) Post-bleach frame in (b) transformed from Cartesian to polar coordinates. (e) Image mask in (c) transformed from Cartesian to polar coordinates. (f) Result of interpolation of data in (d). (g) Model fit of exponential of a Gaussian (Equation (1.1)) (red curve) to median intensity versus distance from bleach-point (in microns) derived from (f) (blue curve).

### Mathematical Methods Supplementary Figure II: FLIP image analysis

Walkthrough of PDE model fitting to FLIP imaging data. (a) Schematic of Partial Differential Equation (PDE) model approximation to FLIP system. (b) Illustration of coarse-grid discretization of cell for numerically solving PDE and fitting to data. (c) Lattice-sites in the grid of (b) are determined as nuclear or cytoplasmic if at least 50% of that lattice site is composed of that cellular compartment. (d) Final coarse-gridded approximation to the original cell. Each grid-point intensity is set at the median intensity of that grid-point. Only pixels from the cellular compartment (nucleus or cytoplasm) that defines the grid-point are used in this calculation. This coarse grid is then used to fit the numerical solution of our PDE to the imaging data.

### Mathematical Methods Supplementary Figure III: FRAP sensitivity analysis

(a) Heatmap of Sum of Squares due to Error (SSE) of the exponential of a Gaussian fit (Equation (1.1)) to the median intensity (Mathematical Methods Supplementary Figure Ig) for varying bleach-depth and effective radius. The red dot signifies the global minimum. (b) Plot of SSE as the bleach-depth parameter is varied, when deriving the bleach-depth parameter from the recovery curve (Equation (1.2)) as opposed to the post-bleach profile. (c) Heatmap of SSE of the single reaction model (Equation (1.5)) fit to the recovery curve of a single cell for varying amplitude and rate of dissociation.

### Mathematical Methods Supplementary Figure IV: FLIP model sensitivity analysis

Sensitivity analysis of model parameters for a single NF1 EYFP-YAP1 WT cell (Plots a-d) and a single CAF1 EYFP-YAP1 WT cell (e-h). (a and e) Plots of sensitivity of association rate, export, import, predicted initial cytoplasmic concentration and decay rate due to bleaching as the fixed dissociation rate is varied. The scales are in terms of ratios of the derived parameter values for a given dissociation rate to the optimum model fit parameter values. (b and f) Plots of sensitivity of association rate, export, import, predicted initial cytoplasmic concentration and decay rate due to bleaching as the fixed rate of diffusion is varied. The scales are in terms of ratios of the derived parameter values for a given rate of diffusion to the optimum model fit parameter values. (c and g) Surface plots of the weighted-SSE (Equation (1.30)) for varying association rate and export. Scales are presented in terms of ratio of parameter values to optimum fit parameter values and ratio of weighted-SSE for those parameter values to the weighted-SSE for optimum parameter values. (d and h) Heatmaps of the weighted-SSE and the overall best-fit weighted-SSE for varying import and export. Scales are again in terms of ratios of specific parameter values to optimum fit parameter values. (i) Horizontal linescan of complete cell for FLIP mathematical solution in Figure S4H at timepoints 0s (blue), 8s (red), 39s (green), 120s (magenta) and 300s (cyan). The discontinuous jumps in intensity illustrate the boundary of the cytoplasm and nucleus.

### Mathematical Methods Supplementary Figure V: FRAP and FLIP residual analysis

Residuals (observed minus predicted values) versus time for FRAP recovery curve model fitting of 30 NF1 EYFP-YAP1 WT cells (ten cells with each of small, medium and large nominal bleach regions) for: (a) pure/effective diffusion model, (b) reaction-diffusion model, (c) single reaction model and (d) double reaction model. Figures e-h constitute a residual analysis of FLIP model fitting to CAF1 EYFP-YAP1 WT cells. (e) Spatial residuals at timepoints 0s, 8s, 39s, 120s and 300s for FLIP model-fitting of a CAFWT cell (see Supplementary Figure S4H). (f) Residuals versus time of FLIP model fit at the nuclear bleachpoint of 15 CAFWT cells. (g) Residuals versus time of FLIP model fit at all nuclear lattice-sites (ignoring bleach-points) for 15 CAFWT cells. (h) Residuals versus time of FLIP model fit at all cytoplasmic lattice-sites of 15 CAFWT cells.

---

**Figure I:**
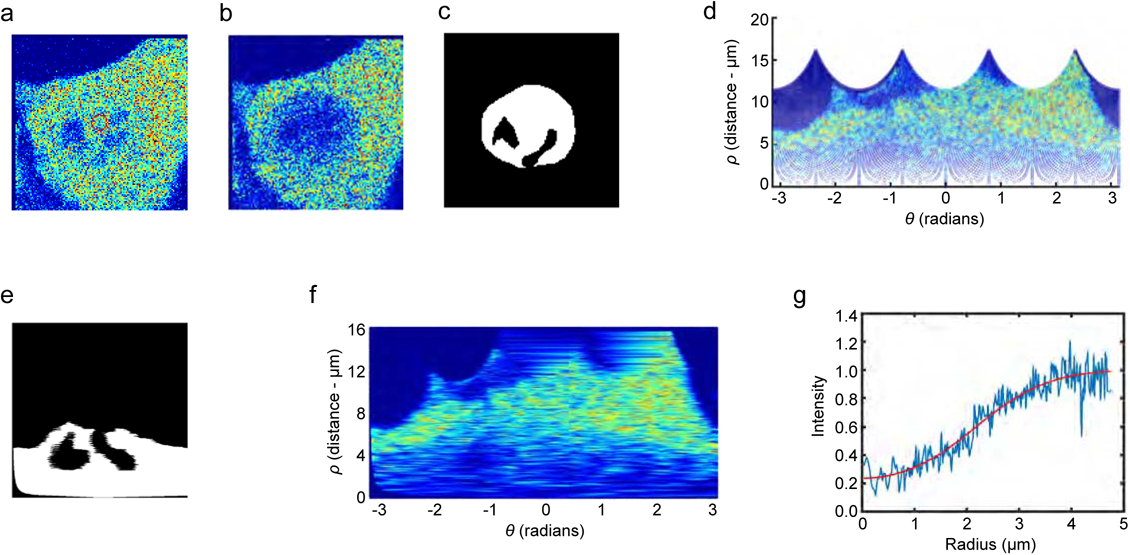
FRAP Image Analysis.

**Figure II:**
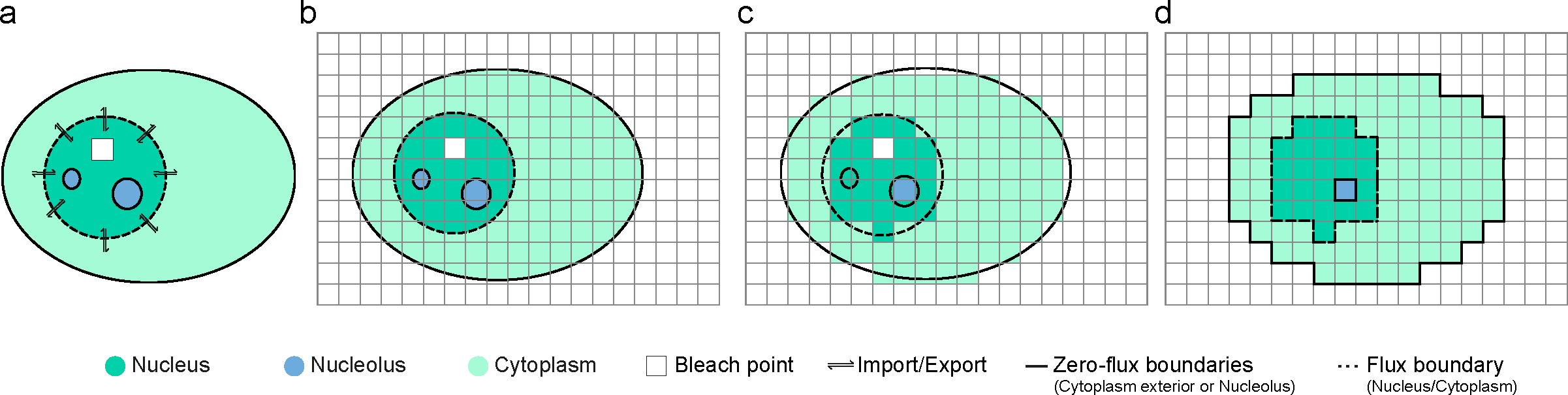
FLIP Image Analysis.

**Figure III:**
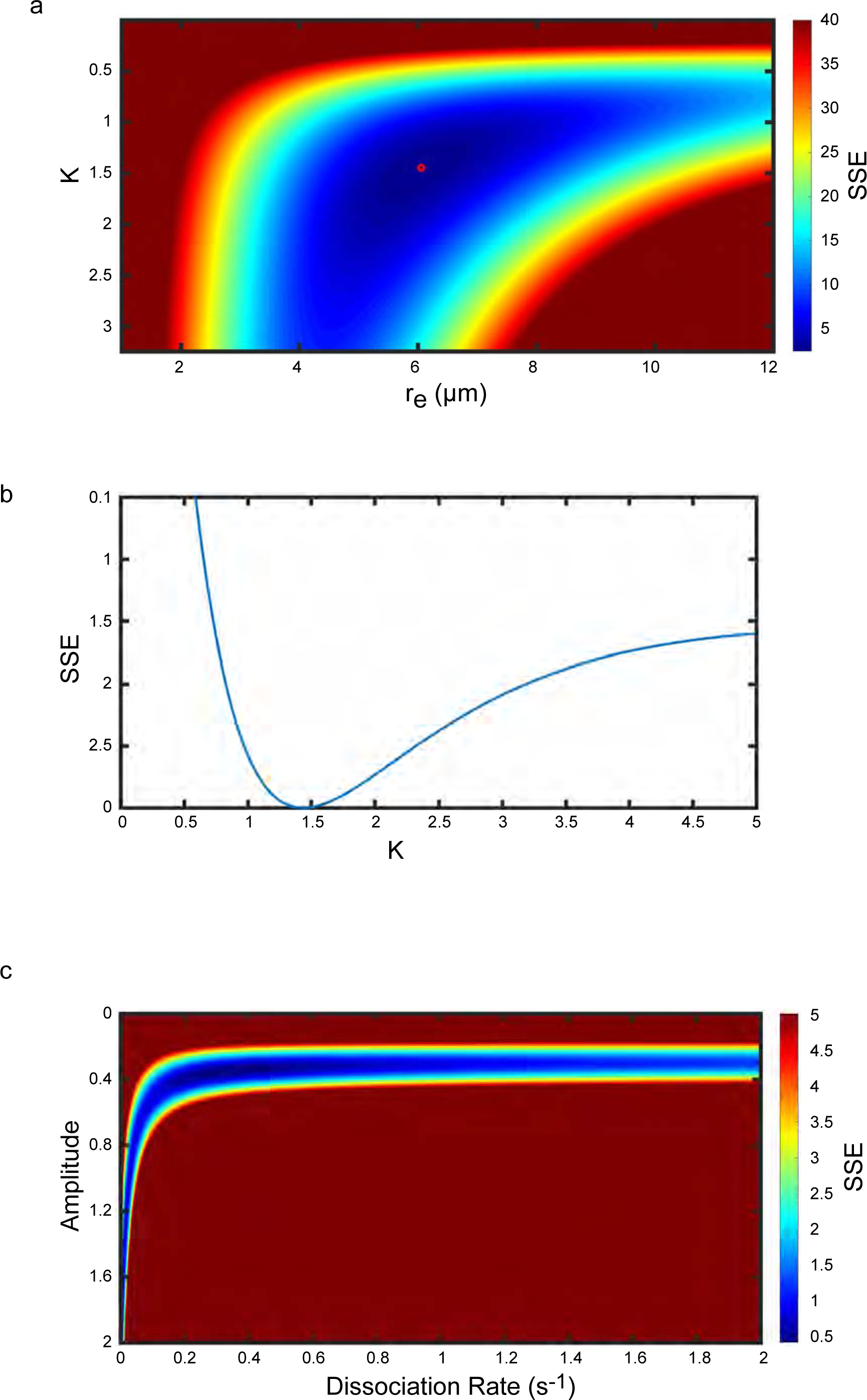
FRAP Sensitivity Analysis.

**Figure IV:**
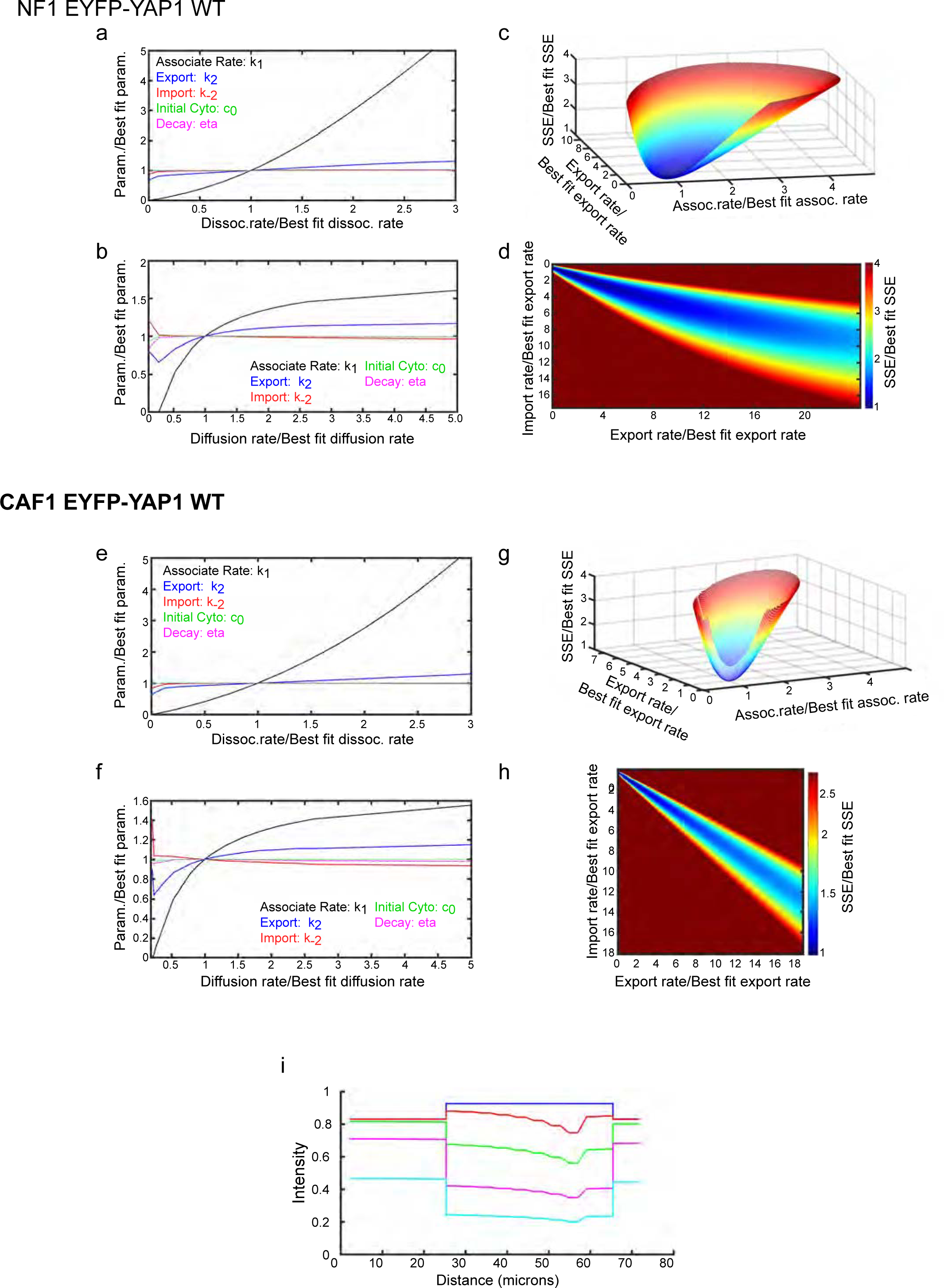
FLIP Sensitivity Analysis -1.

**Figure V:**
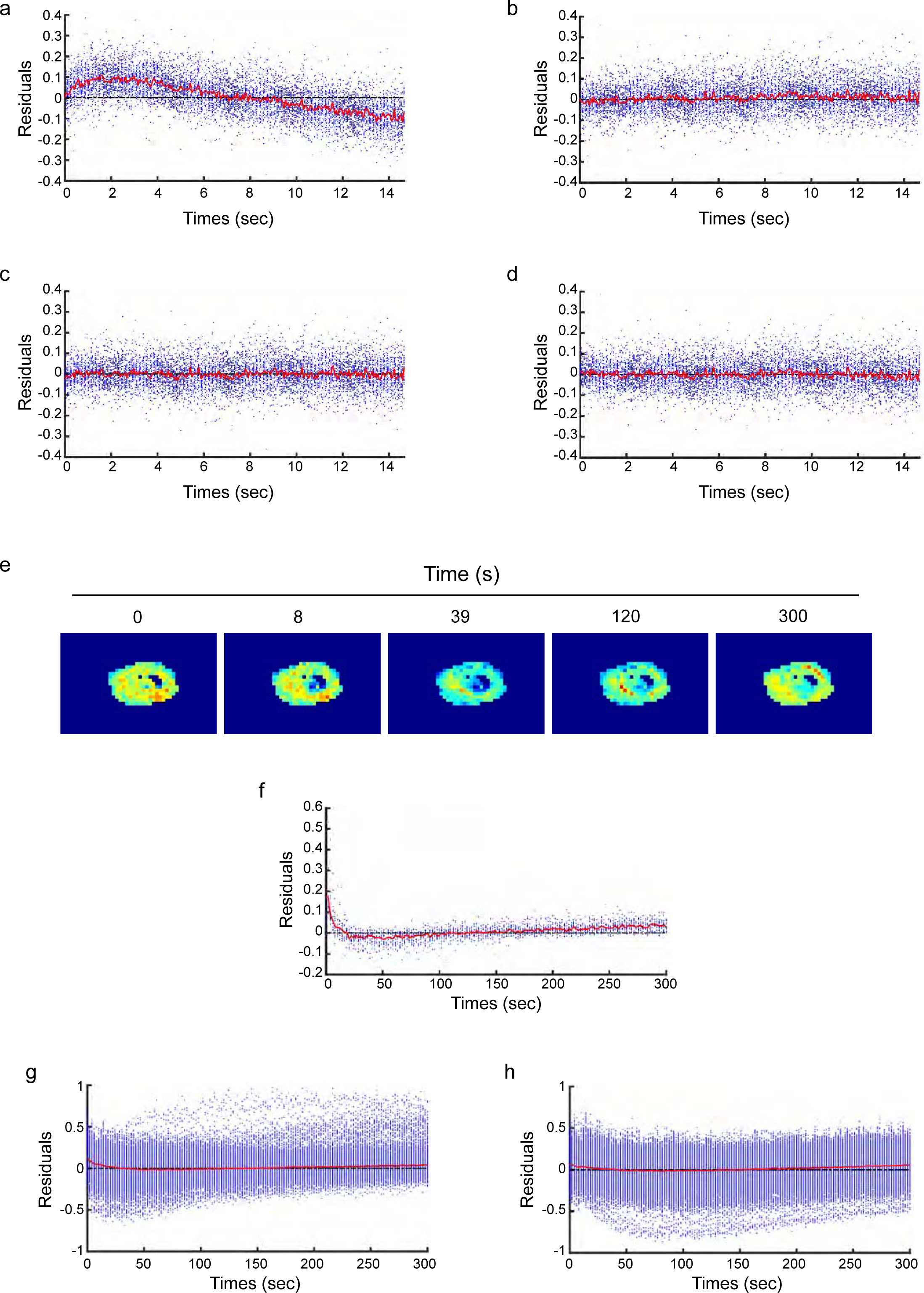
FLIP Sensitivity Analysis-2.

